# Human Endogenous Retrovirus K RNA triggers cytoplasmic TDP-43 aggregation by initiating pathological nuclear condensates

**DOI:** 10.64898/2026.11.16.767080

**Authors:** Yung-Heng Chang, Josh Dubnau

## Abstract

Cytoplasmic aggregation of the nuclear RNA-binding protein TDP-43 is a defining pathology of amyotrophic lateral sclerosis (ALS) and frontotemporal dementia (FTD) and is seen in many cases of Alzheimer’s disease (AD), yet what initiates this process remains unknown. Here we identify an unexpected trigger: RNA from human endogenous retrovirus-K (HERV-K), an ancient retroviral element embedded in the human genome. Using CRISPR-engineered cells expressing Halo-tagged TDP-43 at endogenous levels, we show that HERV-K activation drives formation of cytoplasmic TDP-43 aggregates and functional loss of nuclear TDP-43, measured by aberrant cryptic exon inclusion. Surprisingly, cytoplasmic aggregation is preceded by the appearance of phosphorylated TDP-43 condensates in the nucleus, structures we term Pathological ERV-TDP-43-Associated Nuclear Spheres (PETANS). Live-cell imaging reveals that cytoplasmic inclusions originate directly from these nuclear bodies. This inverts the conventional model to suggest that pathological TDP-43 condensation begins in the nucleus, not the cytoplasm. Super-resolution imaging and structure-function mapping show that HERV-K RNA localizes within PETANS and drives their formation, with a minimal UG-rich non-coding element sufficient to induce the full cascade of nuclear and cytoplasmic pathology. Our findings establish an endogenous retroviral RNA as a direct molecular nucleator of TDP-43 proteinopathy.

## Main

Cells process their biochemical activities within specialized membraneless compartments known as biomolecular condensates whose properties and functions are determined by the specific nucleic acids and proteins contained within. The key role of these biomolecular condensates is to create a microenvironment to facilitate interactions that are essential to conduct cellular reactions^1, 2^. But many proteins that form such biomolecular condensates also are aggregation prone and under dysfunctional conditions contribute to age-dependent neurodegenerative disorders^1^. These include the TDP-43 DNA/RNA binding protein whose proteinopathy is at the core of a suite of disorders that include ALS, FTD, AD and Limbic-predominant Age-related TDP-43 Encephalopathy (LATE)^3, 4, 5^. Under physiological conditions, TDP-43 predominantly localizes to the nucleus where it plays key roles in RNA metabolism. These nuclear roles of TDP-43 include modulating splicing of many cellular mRNAs^6^ and transcriptional silencing of retrotransposons (RTEs) and endogenous retroviruses (ERVs)^7^. In patients with these neurodegenerative diseases, however, TDP-43 protein is seen to mis-localize in both neurons and glial cells to form cytoplasmic inclusions^3, 5, 8^. Many studies also have documented that TDP-43 is at least transiently seen in pathologically hyperphosphorylated form in the nucleus under diverse disease-relevant conditions^9, 10, 11, 12, 13^. Although there are rare disease-causing amino acid substitutions in TDP-43, most cases with TDP-43 proteinopathy involve mislocalization and aggregation of TDP-43 with normal amino acid sequence. The key molecular mechanisms that trigger loss of nuclear localization and deposition of cytoplasmic TDP-43 inclusions are thus important to decipher.

We and others have previously demonstrated that inducing TDP-43 pathology via over-expression or loss of function by knockdown is sufficient to cause the active expression of ERVs and RTEs in both cell and animal models^9, 14, 15, 16, 17, 18^. In the fly model, mdg4 ERV expression causally contributes to the neurodegenerative phenotypes of TDP-43, including its intercellular spread^9, 14, 15^. Increased RTE/ERV expression is also seen in postmortem tissues from ALS and FTD patients^17, 19, 20, 21, 22, 23, 24^. But while ERV expression is caused by loss of TDP-43 nuclear function, we also have previously established that expressing either the fly mdg4-ERV or the human ERV, HERV-K (also known as HML-2), is sufficient to trigger TDP-43 protein pathology in cell models^9^. This suggests the possibility of a feedback amplification in which ERVs somehow cause or exacerbate formation of cytoplasmic inclusions of TDP-43. It remains unknown whether the effects of ERV expression on TDP-43 pathology are direct vs indirect or what the mechanisms underlying this effect might be.

We investigated the mechanisms by which HERV-K impacts TDP-43 protein dysfunction and discovered a key nuclear initiating mechanism. We demonstrate that HERV-K drives nuclear TDP-43 condensates enriched for both HERV-K RNA and pathologically phosphorylated TDP-43 (pTDP-43), which we term Pathological ERV-TDP-43-Associated Nuclear Spheres (PETANS). HERV-K-induced PETANS are initiated in the nucleus and later exit the nucleus to accumulate in the cytoplasm as pathological TDP-43 aggregates that also contain HERV-K RNA. We define a minimal UG-repeat containing RNA region in the HERV-K long terminal repeat (LTR) that induces nuclear PETANS and promotes cytosolic TDP-43 aggregation. These findings support a model in which HERV-K-derived RNA initially drives TDP-43 condensation within the nucleus and subsequently promotes its pathological aggregation in the cytoplasm.

## Results

### HERV-K expression triggers nuclear PETANS and cytoplasmic TDP-43 aggregation

We set out to investigate the mechanisms by which HERV-K expression induces formation of cytoplasmic inclusions of TDP-43. We first used a CRISPR-engineered Halo-tagged TDP-43 cell line^25^, in which endogenous TDP-43 is expressed at physiological levels and is traceable with fluorescent imaging in fixed or live cells. We expressed a series of HERV-K constructs to investigate the structure-to-function relationship between virally encoded components and TDP-43 proteinopathy. ERVs have genetic structure and life cycle that is evolutionarily related to that of exogenous retroviruses (XRVs) such as HIV-1. Replication of ERVs and XRVs share key characteristics including the reverse transcription of their genomic RNA to make a cDNA copy which then is integrated into the host genome. This process utilizes both virally encoded and host encoded components. As with XRVs, ERV first strand cDNA production relies on binding of a host tRNA to the primer binding site (PBS) within the viral mRNA to initiate the process (tRNA-Lys in the case of HERV-K)^26, 27^.

We first tested whether cDNA synthesis is an essential step to trigger TDP-43 pathology. To do so, we generated an H2B-mCherry tagged variant of HERV-K in which the essential cis acting PBS sequence is deleted (HERV-K^ΔPBS^ in Fig. 1a). This engineered HERV-K^ΔPBS^ can produce viral RNA and express each of the virally encoded proteins, but cannot initiate first strand cDNA synthesis. The wild-type H2B-mCherry tagged HERV-K (HERV-K in Fig. 1a) and H2B-mCherry alone (Fig. 1a) were used as positive and negative controls. Each of these three constructs were transfected into the HEK293T cells with endogenously Halo-tagged TDP-43^25^. We then used confocal microscopy to detect the Halo-tagged endogenous TDP-43 and phosphorylated TDP-43 (using an established pTDP-43 antibody, see methods), a marker of pathological TDP-43 inclusions. Imaging was performed at 72 and 120 hours after transfection (Fig. 1b). Within 72-hours after transfection of HERV-K, we saw robust accumulation of cytosolic TDP-43 aggregates. Such cytoplasmic inclusions were detected both with the HaloTag ligand and with the pTDP-43 antibody. These cytoplasmic aggregates remained detectable at 120 hours post transfection (yellow arrowheads in Fig. 1c-e). In contrast, cells expressing H2B-mCherry alone showed almost no detectable cytoplasmic TDP-43 pathology (Fig. 1c-e). We noted that with the HERV-K^ΔPBS^ construct, we saw a significantly higher proportion of cells exhibiting cytosolic TDP-43 aggregates compared with wild-type HERV-K expression (yellow arrowheads in Fig. 1c-e). Although we do not understand the reasons why HERV-K^ΔPBS^ drives even higher levels of TDP-43 pathology, the differential effect between HERV-K and HERV-K^ΔPBS^ was not attributable to differences in expression levels between constructs (Extended Data Fig. 1a-c) or to effects on endogenous TDP-43 abundance (Extended Data Fig. 1d). In addition to the effects on cytoplasmic TDP-43 aggregates, the expression of either wild-type HERV-K or HERV-K^ΔPBS^ constructs caused the appearance of spherical nuclear structures that we designated as Pathological ERV-TDP-43-Associated Nuclear Spheres (PETANS) (pink arrowheads in Fig. 1c-e and Extended Data Fig. 1e,f). The outer surface of PETANS was labeled with both the HaloTag ligand and the pTDP-43 antibody indicating that PETANS contained pathological TDP-43 condensates (Fig. 1c and Extended Data Fig. 1e,f) exhibiting a broad and variable size range (Extended Data Fig. 1g).

**Fig 1.**
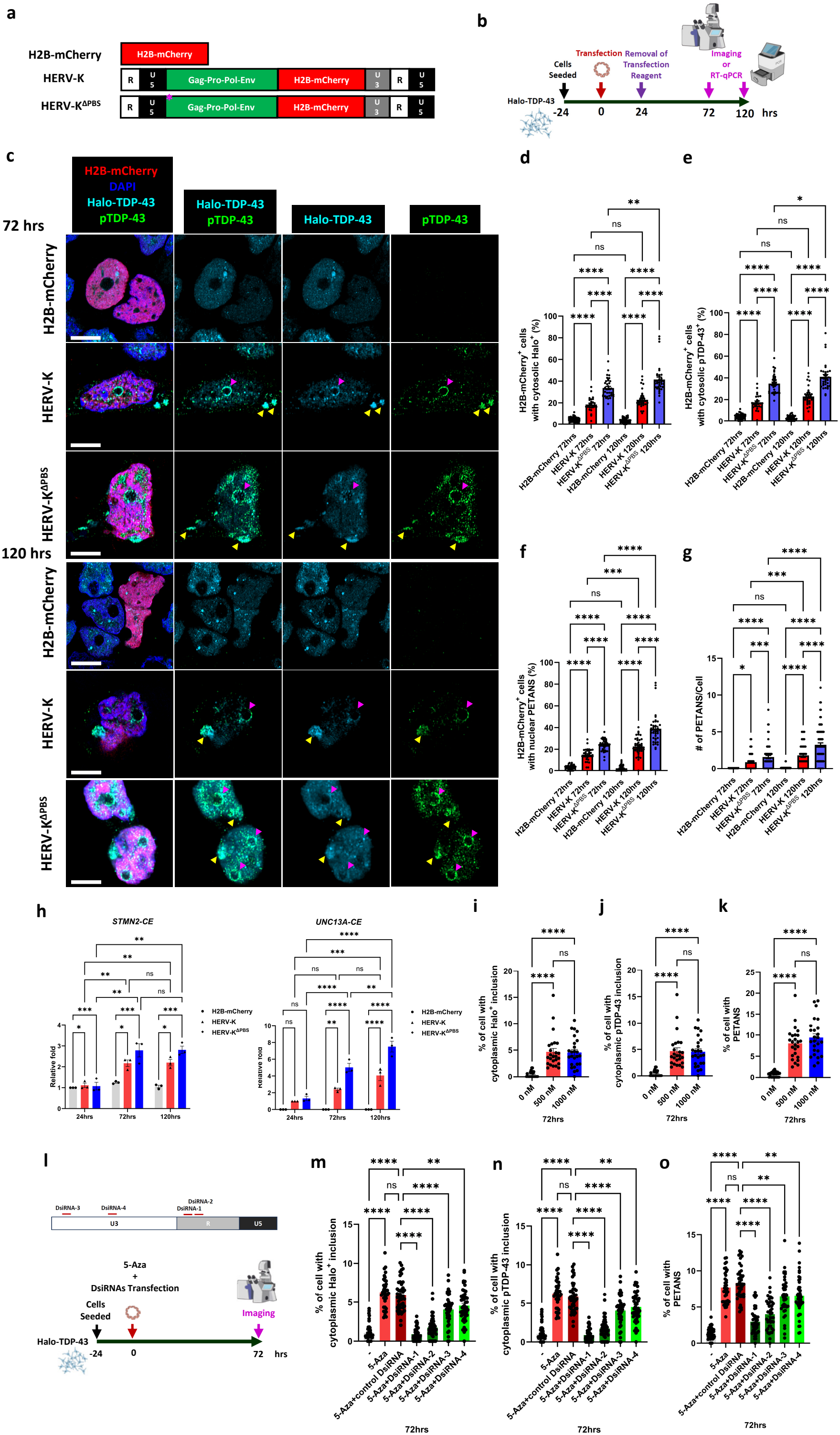
HERV-K expression triggers nuclear PETANS, cytosolic TDP-43 aggregates, and cryptic exon inclusion of TDP-43 target RNA. **a**, Schematic description of control H2B-mCherry, HERV-K (HERV-K-H2B-mCherry) and HERV-K^ΔPBS^ (HERV-K^ΔPBS^-H2B-mCherry) constructs. **b**, Illustration of the experimental timeframe for transfection conditions and visualization. **c**, Subcellular localization of nuclear PETANS (pink arrowhead) and cytosolic TDP-43 aggregates (yellow arrowhead) were visualized by Halo-TDP-43 (cyan), pTDP-43 (green), H2B-mCherry (red), and DAPI (blue) in response to HERV-K and HERV-K^ΔPBS^ expression. Scale bar=5 µm. Quantification of HERV-K and HERV-K^ΔPBS^ transfected cells showing cytoplasmic **d**, Halo-TDP-43 (72 hours: H2B-mCherry 4.66±0.28%, n=56; HERV-K 18.52±1.24%, n=27 and HERV-K^ΔPBS^ 33.72±1.47%, n=35. 120 hours: H2B-mCherry 3.89±0.41%, n=35; HERV-K 22.89±1.55%, n=33 and HERV-K^ΔPBS^ 41.52±2.31%, n=34) and **e**, pTDP-43 (72 hours: H2B-mCherry 4.31±0.32%, n=56; HERV-K 17.50±1.32%, n=27 and HERV-K^ΔPBS^ 34.76±1.49%, n=35. 120 hours: H2B-mCherry 3.08±0.37%, n=35; HERV-K 22.98±1.57%, n=33 and HERV-K^ΔPBS^ 41.08±2.11%, n=34) aggregation. Data are shown as mean ± SEM and a two-way ANOVA with multiple comparisons was performed. *p<0.05; **p<0.01; ****p<0.0001; ns, no significant difference. **f**, Quantification of transfected cells developing nuclear PETANS after HERV-K and HERV-K^ΔPBS^ expression (72 hours: H2B-mCherry 3.24±0.27%, n=56; HERV-K 15.14±0.93%, n=34 and HERV-K^ΔPBS^ 23.94±0.79%, n=43. 120 hours: H2B-mCherry 2.57±0.47%, n=35; HERV-K 22.63±1.21%, n=42 and HERV-K^ΔPBS^ 38.96±2.19%, n=40). **g**, PETANS number per-transfected cell (72 hours: H2B-mCherry 0.00±0.00, n=90 PETANS; HERV-K 0.88±0.11, n=102 PETANS and HERV-K^ΔPBS^ 1.58±0.23, n=59 PETANS. 120 hours: H2B-mCherry 0.04±0.03, n=50 PETANS; HERV-K 1.80±0.27, n=35 PETANS and HERV-K^ΔPBS^ 3.24±0.26, n=83 PETANS) after HERV-K and HERV-K^ΔPBS^ expression. Data are shown as mean ± SEM and a two-way ANOVA with multiple comparisons was performed. *p<0.05; ***p<0.001; ****p<0.0001; ns, no significant difference. **h**, RT-qPCR analysis was performed to validate the expression of *STMN2-CE* and *UNC13A-CE* inclusions after HERV-K and HERV-K^ΔPBS^ expression. Data are shown as mean ± SEM with n=3 biological replicates per timepoint and a two-way ANOVA with multiple comparisons was performed. *p<0.05; **p<0.01; ****p<0.0001; ns, no significant difference. Halo-TDP-43 cells developed **i**, cytosolic Halo-TDP-43 aggregates (0 nM 0.37±0.11%, n=25; 500 nM 4.67±0.64%, n=25 and 1000 nM 4.56±0.56%, n=25), **j**, cytosolic pTDP-43 aggregates (0 nM 0.37±0.11%, n=25; 500 nM 4.67±0.64%, n=25 and 1000 nM 4.56±0.56%, n=25), and **k**, nuclear PETANS (0 nM 0.68±0.12%, n=25; 500 nM 8.00±0.75%, n=25 and 1000 nM 9.41±0.87%, n=25) by 5-Aza treatment after 72 hours treatment. Data are shown as mean ± SEM and a one-way ANOVA with multiple comparisons was performed. ****p<0.0001; ns, no significant difference. **l**, Schematic description of HERV-K DsiRNA target sites and the experimental timeframe for transfection conditions and 5-Aza treatment. Halo-TDP-43 cells developed **m**, cytosolic Halo-TDP-43 aggregates (−1.06±0.14%, n=52; 5-Aza 6.28±0.30%, n=39; 5-Aza+control DsiRNA 5.90±0.30%, n=44; 5-Aza+DsiRNA-1 0.89±0.13%, n=43; 5-Aza+DsiRNA-2 1.74±0.21%, n=42; 5-Aza+DsiRNA-3 4.05±0.30%, n=41; 5-Aza+DsiRNA-4 4.56±0.29%, n=47), **n**, cytosolic pTDP-43 aggregates (−1.06±0.14%, n=52; 5-Aza 6.28±0.30%, n=39; 5-Aza+control DsiRNA 5.84±0.29%, n=44; 5-Aza+DsiRNA-1 0.89±0.13%, n=43; 5-Aza+DsiRNA-2 1.74±0.21%, n=42; 5-Aza+DsiRNA-3 4.05±0.30%, n=41; 5-Aza+DsiRNA-4 4.52±0.29%, n=47), and **o**, nuclear PETANS (−1.31±0.10%, n=52; 5-Aza 7.68±0.35%, n=39; 5-Aza+control DsiRNA 8.31±0.34%, n=44; 5-Aza+DsiRNA-1 2.97±0.26%, n=43; 5-Aza+DsiRNA-2 4.01±0.32%, n=42; 5-Aza+DsiRNA-3 6.46±0.41%, n=41; 5-Aza+DsiRNA-4 6.56±0.38%, n=47) following 72 hours of 5-Aza and HERV-K DsiRNAs treatment. Data is shown as mean ± SEM and a one-way ANOVA with multiple comparisons was performed. **p<0.01; ****p<0.0001; ns, no significant difference.

As with the cytoplasmic aggregates, we saw accumulation of PETANS in the nucleus with expression of either the wild-type HERV-K or the HERV-K^ΔPBS^. PETANS were observed at 72 hours after HERV-K expression, with increased numbers at 120 hours (Fig. 1c,f,g). Cytoplasmic inclusions that were uniformly labeled with both Halo-tagged and phosphorylated TDP-43 were seen after expression of either the HERV-K or the HERV-K^ΔPBS^ constructs. But in HERV-K^ΔPBS^-expressing cells, we also occasionally detected spherical PETANS like structures with Halo-tag and pTDP-43 labeling present in the cytoplasm (Fig. 1c). The observed pathological effects on TDP-43 localization after HERV-K and HERV-K^ΔPBS^ expression also were associated with functional effects on inclusion of cryptic exons (CEs) in well-established target RNAs, *STMN2*^*28, 29*^ and *UNC13A*^*30, 31*^. RT-qPCR analysis revealed that CE inclusion in the *STMN2* and *UNC13A* transcripts were gradually increased over time in HERV-K overexpressing cells (Fig. 1h). This effect was exacerbated in the presence of the HERV-K^ΔPBS^ compared with that of wild-type HERV-K (Fig. 1h). Together, these results indicate HERV-K overexpression can promote the formation of both spherical nuclear condensates with pTDP-43 on the surface and cytosolic aggregates of TDP-43, while concurrently disrupting the regulation of canonical TDP-43 target RNAs, *STMN2* and *UNC13A*. Importantly, deletion of the PBS, a cis-acting element essential for cDNA synthesis, does not eliminate the pathological effects of HERV-K expression on the formation of PETANS, of cytoplasmic TDP-43 inclusions or of CE inclusions in *STMN2* and *UNC13A*. These findings support the notion that cDNA synthesis is not essential to the mechanism by which HERV-K triggers TDP-43 pathology.

We also investigated whether HERV-K affects other ALS-associated, predominantly nuclear ribonucleoproteins, specifically Fused in Sarcoma (FUS) and Heterogeneous Nuclear Ribonucleoprotein A1 (hnRNP A1)^32, 33, 34^, and whether these proteins associate with PETANS.

Immunostaining revealed that neither FUS nor hnRNP A1 colocalized with nuclear PETANS in HERV-K^ΔPBS^ expressing cells (Extended Data Fig. 2a,b). Among HERV-K^ΔPBS^-expressing cells exhibiting cytoplasmic TDP-43 pathology, only a small proportion showed cytosolic co-localization of FUS with TDP-43, whereas virtually no hnRNP A1 co-localized with TDP-43 was observed (Extended Data Fig. 2a,b). These findings suggest that HERV-K preferentially promotes TDP-43 proteinopathy rather than comparable ALS-associated FUS or hnRNP A1 aggregation.

**Fig 2.**
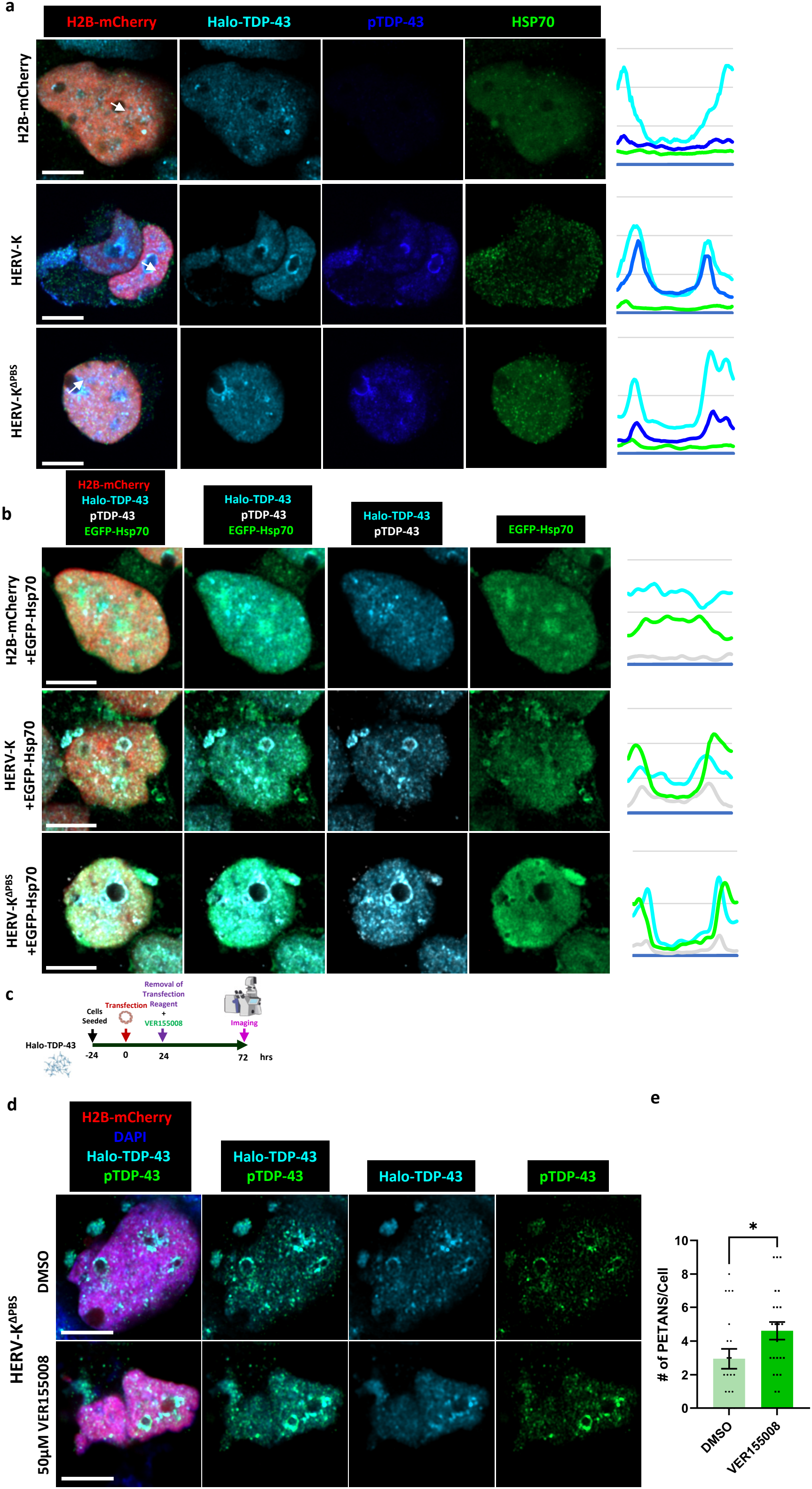
Hsp70 inhibitor treatment did not disrupt PETANS formation. **a**, Visualization of the subcellular localization of endogenous Hsp70 protein in Halo-TDP-43 cells after HERV-K and HERV-K^ΔPBS^ expression, co-stained with Hsp70 (green), Halo-TDP-43 (cyan), pTDP-43 (blue), and H2B-mCherry (red). Scale bar=5 µm. **b**, Visualization of the subcellular localization of ectopic EGFP-Hsp70 protein in Halo-TDP-43 cells after HERV-K and HERV-K^ΔPBS^ expression, co-stained with EGFP-Hsp70 (green), Halo-TDP-43 (cyan), pTDP-43 (white), and H2B-mCherry (red). Scale bar=5 µm. **c**, Schematic description of the experimental timeframe for HERV-K^ΔPBS^ transfection conditions and Hsp70 inhibitor VER155008 treatment. **d**, Visualization of the effect of VER155008 treatment on HERV-K^ΔPBS^-induced PETANS in Halo-TDP-43 cells, co-stained with Halo-TDP-43 (cyan), pTDP-43 (green), H2B-mCherry (red), and DAPI (blue). Scale bar=5 µm. **e**, Quantification of nuclear PETANS number in HERV-K^ΔPBS^ transfected cells with 50 µM VER155008 treatment (DMSO 2.95±0.59, n=20 PETANS and VER155008 4.61±0.52, n=23 PETANS). Data are shown as mean ± SEM and an unpaired t-test was performed. *p<0.05.

We previously demonstrate that expression of *Drosophila* mdg4-ERV in *Drosophila* S2 cells also promotes cytosolic TDP-43 aggregation^9^. As with HERV-K, we tested whether deleting the PBS from mdg4-ERV had an impact on the mdg4-ERV’s ability to trigger TDP-43 pathology. We co-transfected wild-type or PBS-deleted mdg4-ERV constructs along with an inducible human TDP-43 tagged with V5 (TDP-43-V5) for 72 hours (Extended Data Fig. 3a,b) in *Drosophila* S2 cells. Following induction of hTDP-43-V5 expression for 20 hours, cells co-expressing either form of mdg4-ERV displayed robust cytoplasmic aggregation of phosphorylated hTDP-43 (Extended Data Fig. 3c-e). These results parallel the effects observed with HERV-K, suggesting that the molecular mechanisms by which each of these ERVs promote TDP-43 pathology is conserved and is independent of cDNA production and therefore is most likely due to some combination of the RNA transcripts and/or the encoded protein products.

**Fig 3.**
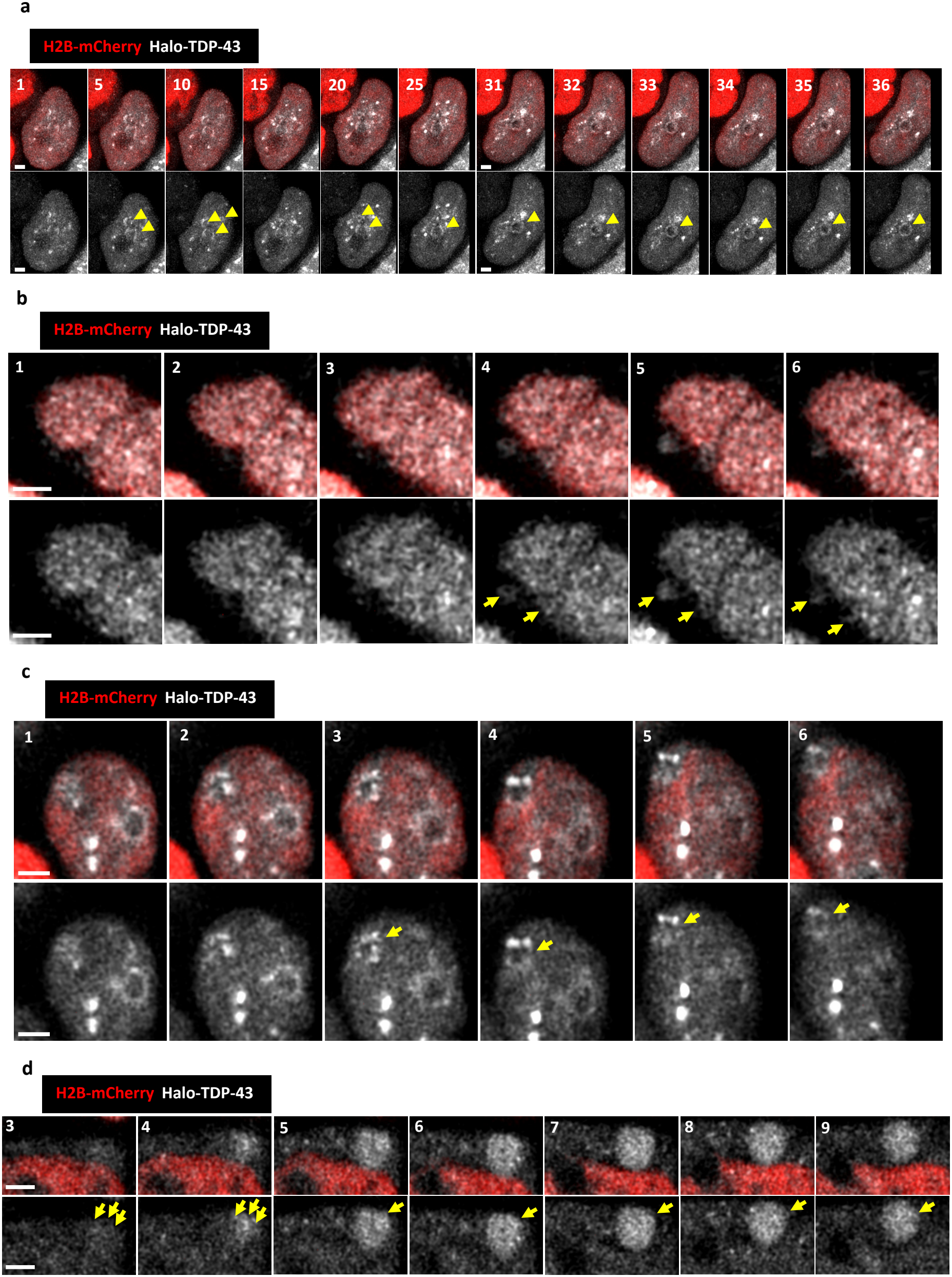
Live-cell imaging of PETANS assembly in the nucleus, secretion from the nucleus and assembly of aggregates in the cytoplasm. **a**, A series of images captured from Extended Data Movie 1, showing PETANS assembly with staining for Halo-TDP-43 (white) and H2B-mCherry (red). Scale bar=2 µm. **b**, and **c**, A series of images captured from Extended Data Movie 2 and Extended Data Movie 3, showing PETANS release from the nucleus with staining for Halo-TDP-43 (white) and H2B-mCherry (red). Scale bar=2 µm. **d**, A series of images captured from Extended Data Movie 4, showing multiple PETANS forming aggregate in the cytoplasm with staining for Halo-TDP-43 (white) and H2B-mCherry (red). Scale bar=2 µm.

### HERV-K induces nuclear PETANS in a variety of human cell lines

We wondered whether the HERV-K-induced PETANS and cytoplasmic aggregates that we observed could be an artifact caused by the expression of a HaloTag, the fusion of the HaloTag to TDP-43 or of the HEK293 cell context. We tested whether PETANS or cytoplasmic inclusions of TDP-43 were present in HERV-K-expressing HEK293T cells where endogenous GAPDH gene, rather than TDP-43, was tagged with HaloTag. In the Halo-GAPDH cells, HERV-K overexpression caused nuclear PETANS and cytosolic TDP-43 aggregates that were detected with the pTDP-43 antibody (Extended Data Fig. 4a), whereas these features were absent in the same Halo-GAPDH cells when transfected with H2B-mCherry control. Importantly, the Halo ligand signal did not colocalize with the pTDP-43-labeled nuclear PETANS or cytosolic TDP-43 aggregates in these Halo-GAPDH HEK293 cells (Extended Data Fig. 4a). Thus, PETANS formation is dependent on the expression of HERV-K, and the HaloTag ligand fluorescence only colocalizes with these features of TDP-43 pathology when the HaloTag is fused to the endogenous TDP-43 encoding gene, not when it is fused to GAPDH. Next, we ectopically expressed HERV-K or HERV-K^ΔPBS^ in wild-type HEK293T cells that had not been engineered to express any Halo-tagged proteins. Both HERV-K and HERV-K^ΔPBS^ induced nuclear PETANS and cytosolic TDP-43 aggregates, which were independently detected using pan-TDP-43 or pTDP-43 antibodies. In contrast, these pathological TDP-43 features were not observed in H2B-mCherry transfected control cells (Extended Data Fig. 4b). Thus, PETANS formation is not an artifact of Halo-tagging of TDP-43 or other endogenous proteins.

**Fig 4.**
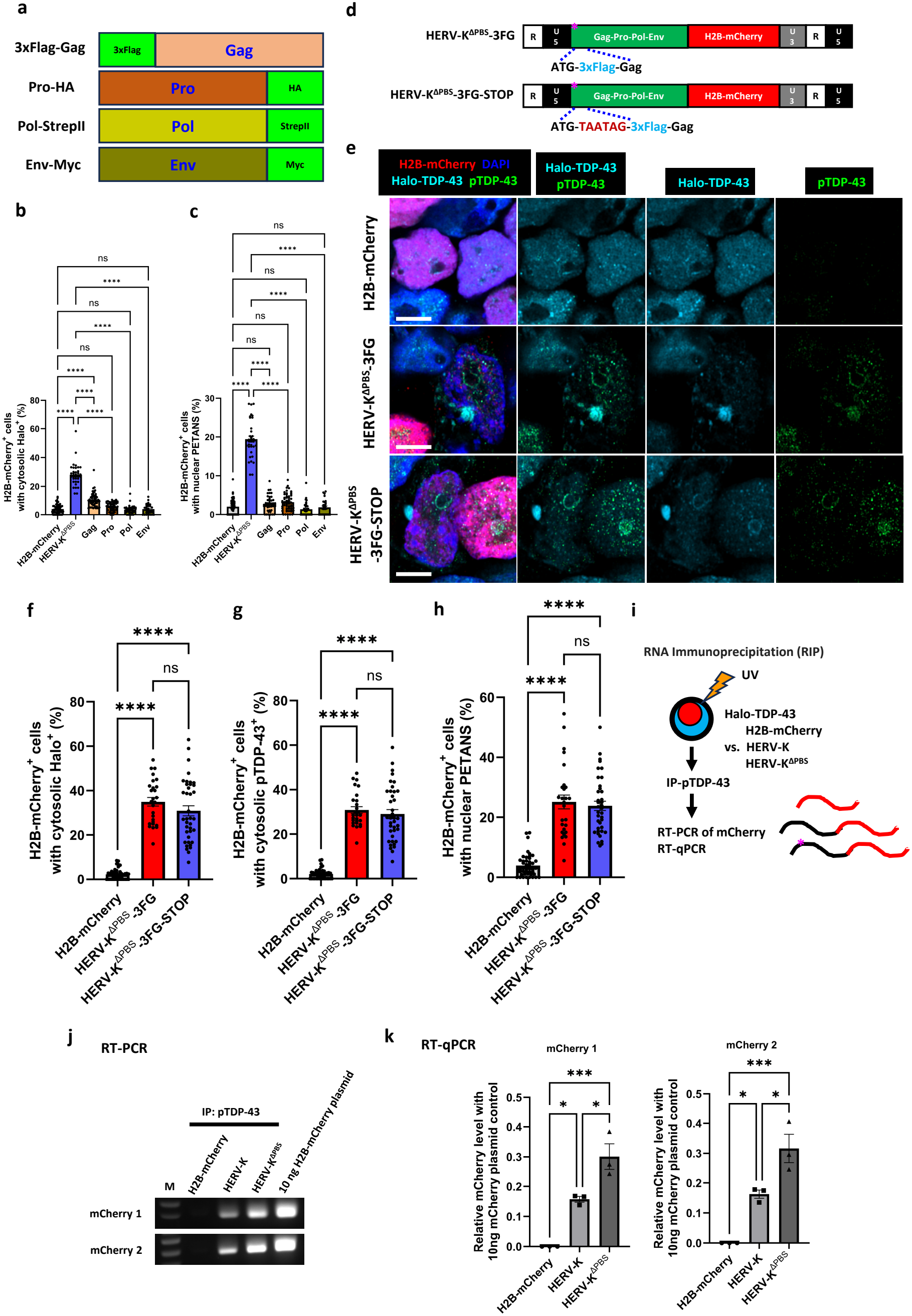
HERV-K RNA, but not the coding ORFs, triggers PETANS and cytosolic TDP-43 aggregates. **a**, Schematic description of HERV-K ORF expressing constructs fused with different tags. Quantification of HERV-K ORF-transfected cells showing **b**, cytoplasmic Halo-TDP-43 (H2B-mCherry 3.96±0.40%, n=61; HERV-K^ΔPBS^ 28.18±1.30%, n=36; Gag 10.48±0.64%, n=57; Pro 5.74±0.40%, n=55; Pol 3.63±0.52%, n=33; Env 3.91±0.61%, n=30) and developing **c**, nuclear PETANS (H2B-mCherry 2.03±0.24%, n=61; HERV-K^ΔPBS^ 19.44±0.79%, n=36; Gag 2.78±0.29%, n=45; Pro 3.18±0.29%, n=55; Pol 1.37±0.31%, n=33; Env 1.87±0.34%, n=30). Data are shown as mean ± SEM and a one-way ANOVA with multiple comparisons was performed. ****p<0.0001; ns, no significant difference. **d**, Schematic description of HERV-K^ΔPBS^-3FG and HERV-K^ΔPBS^-3FG-STOP expressing constructs. **e**, Nuclear PETANS and cytosolic TDP-43 aggregates were visualized by Halo-TDP-43 (cyan), pTDP-43 (green), H2B-mCherry (red), and DAPI (blue) after HERV-K^ΔPBS^-3FG and HERV-K^ΔPBS^-3FG-STOP expression. Scale bar=5 µm. Quantification of control H2B-mCherry, HERV-K^ΔPBS^-3FG and HERV-K^ΔPBS^-3FG-STOP transfected cells showing cytoplasmic **f**, Halo-TDP-43 (H2B-mCherry 2.31±0.34%, n=46; HERV-K^ΔPBS^-3FG 34.98±1.93%, n=27 and HERV-K^ΔPBS^-3FG-STOP 30.90±2.27%, n=38) and **g**, pTDP-43 (H2B-mCherry 2.09±0.32%, n=46; HERV-K^ΔPBS^-3FG 30.84±1.51%, n=27 and HERV-K^ΔPBS^-3FG-STOP 29.00±2.01%, n=38) aggregation. Data are shown as mean ± SEM and a one-way ANOVA with multiple comparisons was performed. ****p<0.0001; ns, no significant difference. **h**, Quantification of transfected cells developing nuclear PETANS after control H2B-mCherry, HERV-K^ΔPBS^-3FG or HERV-K^ΔPBS^-3FG-STOP expression (H2B-mCherry 3.94±0.54%, n=46; HERV-K^ΔPBS^-3FG 25.12±2.32%, n=27 and HERV-K^ΔPBS^-3FG-STOP 23.82±1.56%, n=38). Data are shown as mean ± SEM and a one-way ANOVA with multiple comparisons was performed. ****p<0.0001; ns, no significant difference. **i**, Schematic illustration of the experimental flowchart to characterize the physical interaction between pTDP-43 protein and HERV-K RNAs by RIP. **j**, RT-PCR and **k**, RT-qPCR results of control H2B-mCherry, HERV-K and HERV-K^ΔPBS^ RNAs interacting with pTDP-43 protein. Data are shown as mean ± SEM with n=3 biological replicates per experimental condition and a one-way ANOVA with multiple comparisons was performed. *p<0.05; ***p<0.001.

Next, we examined whether HERV-K expression could cause PETNAS formation in other human cell types. To experimentally test this, we expressed HERV-K or HERV-K^ΔPBS^ in a range of commonly used immortalized cells derived from human microglia (HMC3), astrocytes (SVG p12), and neuroblastoma (SH-SY5Y) cells. In all three cell types, HERV-K or HERV-K^ΔPBS^ overexpression led to cytosolic TDP-43 aggregation, whereas this phenotype was absent in control H2B-mCherry cells (Extended Data Fig. 4c-e). Notably, both HERV-K and HERV-K^ΔPBS^ also triggered nuclear PETANS formation across all examined cell types, which was not observed in H2B-mCherry controls (Extended Data Fig. 4c-e). Collectively, these results demonstrate that HERV-K expression can induce nuclear PETANS and cytoplasmic aggregates of TDP-43 across diverse human cell types.

### Endogenous HERV-K reactivation causes nuclear PETANS and cytosolic TDP-43 aggregates

The above findings derive from expression of HERV-K from transfected HERV-K constructs. We next asked whether activation of endogenous HERV-K would produce similar effects. To achieve this, DNA methyltransferase inhibitor, 5-azacytidine (5-Aza), was used to induce expression of epigenetically silenced ERVs and RTEs, including the HERV-K^35, 36^. Dose-response experiments (0 to 1000 nM) revealed that treatment with 500 or 1000 nM 5-Aza for 72 hours caused significant activation of HERV-K expression (Extended Data Fig. 5a,b). Notably, either concentration of 5-Aza treatment triggered robust nuclear PETANS formation and cytosolic TDP-43 aggregation compared with untreated cells (Fig. 1i-k and Extended Data Fig. 5c). Concurrently, 5-Aza treatment promoted CE inclusion in *STMN2* transcripts (Extended Data Fig. 5d), but did not significantly increase *UNC13A-CE* inclusion (Extended Data Fig. 5d). This difference may reflect the relatively lower level of TDP-43 pathology induced by 5-Aza treatment compared with HERV-K overexpression. Differential responses among TDP-43 target transcripts have been reported previously and this is thought to depend on the severity of TDP-43 dysfunction^37^.

**Fig 5.**
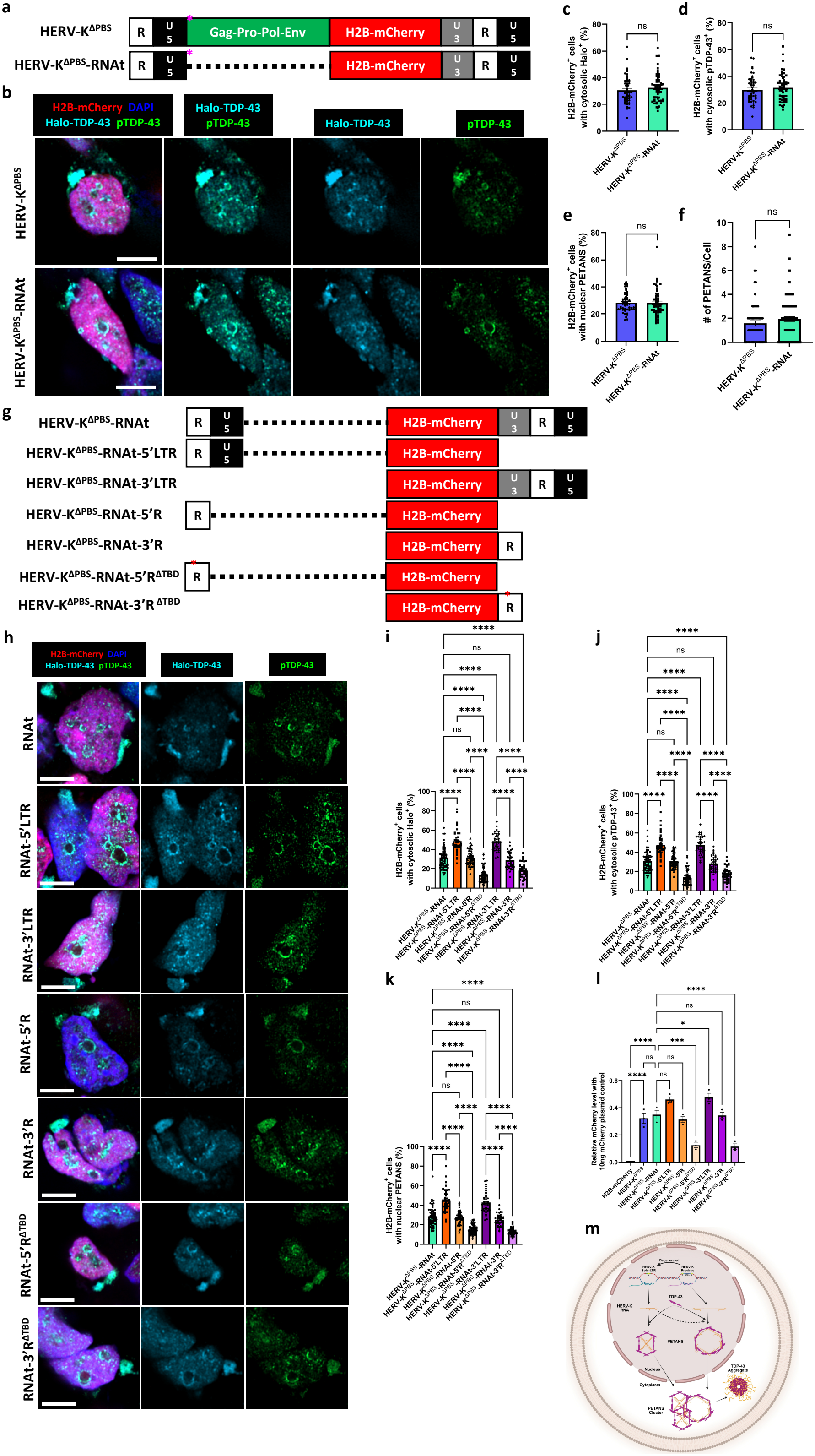
The R element of the HERV-K LTR is sufficient to induce PETANS and cytosolic TDP-43 aggregates. **a**, Schematic description of the HERV-K^ΔPBS^-RNAt expressing construct. **b**, Nuclear PETANS and cytosolic TDP-43 aggregates were visualized by Halo-TDP-43 (cyan), pTDP-43 (green), H2B-mCherry (red), and DAPI (blue) under HERV-K^ΔPBS^ and HERV-K^ΔPBS^-RNAt expression. Scale bar=5 µm. Quantification of HERV-K^ΔPBS^ and HERV-K^ΔPBS^-RNAt transfected cells inducing cytoplasmic **c**, Halo-TDP-43 (HERV-K^ΔPBS^ 30.69±1.41%, n=52 and HERV-K^ΔPBS^-RNAt 32.59±1.44%, n=54) and **d**, pTDP-43 (HERV-K^ΔPBS^ 30.03±1.30%, n=52 and HERV-K^ΔPBS^-RNAt 31.58±1.38%, n=54) aggregation. **e**, Quantification of transfected cells developing nuclear PETANS after HERV-K^ΔPBS^ or HERV-K^ΔPBS^-RNAt expression (HERV-K^ΔPBS^ 28.43±0.96%, n=52 and HERV-K^ΔPBS^-RNAt 28.23±1.40%, n=54). **f**, PETANS number of per-transfected cell (HERV-K^ΔPBS^ 1.58±0.23, n=59 PETANS and HERV-K^ΔPBS^-RNAt 1.94±0.20, n=82 PETANS) after HERV-K^ΔPBS^ or HERV-K^ΔPBS^-RNAt expression. Data are shown as mean ± SEM and unpaired t-test was performed. ns, no significant difference. **g**, Schematic description of the HERV-K^ΔPBS^-RNAt, HERV-K^ΔPBS^-RNAt-5’LTR, HERV-K^ΔPBS^-RNAt-3’LTR, HERV-K^ΔPBS^-RNAt-5’R, HERV-K^ΔPBS^-RNAt-3’R, HERV-K^ΔPBS^-RNAt-5’R^ΔTBD^ and HERV-K^ΔPBS^-RNAt-3’R^ΔTBD^ expressing constructs. **h**, Nuclear PETANS and cytosolic TDP-43 aggregates were visualized by Halo-TDP-43 (cyan), pTDP-43 (green), H2B-mCherry (red), and DAPI (blue) after HERV-K^ΔPBS^-RNAt, HERV-K^ΔPBS^-RNAt-5’LTR, HERV-K^ΔPBS^-RNAt-3’LTR, HERV-K^ΔPBS^-RNAt-5’R, HERV-K^ΔPBS^-RNAt-3’R, HERV-K^ΔPBS^-RNAt-5’R^ΔTBD^ or HERV-K^ΔPBS^-RNAt-3’R^ΔTBD^ expression. Scale bar=5 µm. Quantification of HERV-K^ΔPBS^-RNAt, HERV-K^ΔPBS^-RNAt-5’LTR, HERV-K^ΔPBS^-RNAt-3’LTR, HERV-K^ΔPBS^-RNAt-5’R, HERV-K^ΔPBS^-RNAt-3’R, HERV-K^ΔPBS^-RNAt-5’R^ΔTBD^ or HERV-K^ΔPBS^-RNAt-3’R^ΔTBD^ transfected cells triggering cytoplasmic **i**, Halo-TDP-43 (HERV-K^ΔPBS^-RNAt 31.45±1.38%, n=60; HERV-K^ΔPBS^-RNAt-5’LTR 48.73±1.87%, n=40; HERV-K^ΔPBS^-RNAt-5’R 31.22±1.06%, n=53; HERV-K^ΔPBS^-RNAt-5’R^ΔTBD^ 13.24±0.98%, n=59; HERV-K^ΔPBS^-RNAt-3’LTR 48.73±1.38%, n=40; HERV-K^ΔPBS^-RNAt-3’R 28.62±1.32%, n=40 and HERV-K^ΔPBS^-RNAt-3’R^ΔTBD^ 18.06±1.16%, n=40) and **j**, pTDP-43 (HERV-K^ΔPBS^-RNAt 30.54±1.32%, n=60; HERV-K^ΔPBS^-RNAt-5’LTR 47.75±1.79%, n=40; HERV-K^ΔPBS^-RNAt-5’R 30.76±1.01%, n=53; HERV-K^ΔPBS^-RNAt-5’R^ΔTBD^ 13.12±0.96%, n=59; HERV-K^ΔPBS^-RNAt-3’LTR 47.69±1.42%, n=40; HERV-K^ΔPBS^-RNAt-3’R 28.13±1.24%, n=40 and HERV-K^ΔPBS^-RNAt-3’R^ΔTBD^ 17.41±1.08%, n=40) aggregation. **k**, Quantification of transfected cells developing nuclear PETANS after HERV-K^ΔPBS^ or HERV-K^ΔPBS^-RNAt expression (HERV-K^ΔPBS^-RNAt 28.43±1.26%, n=60; HERV-K^ΔPBS^-RNAt-5’LTR 45.12±1.84%, n=40; HERV-K^ΔPBS^-RNAt-5’R 26.97±0.88%, n=53; HERV-K^ΔPBS^-RNAt-5’R^ΔTBD^ 15.05±0.58%, n=59; HERV-K^ΔPBS^-RNAt-3’LTR 42.95±1.60%, n=40; HERV-K^ΔPBS^-RNAt-3’R 24.95±0.93%, n=40 and HERV-K^ΔPBS^-RNAt-3’R^ΔTBD^ 12.83±0.57%, n=40). Data are shown as mean ± SEM and a one-way ANOVA with multiple comparisons was performed. ****p<0.0001; ns, no significant difference. **l**, Using RT-qPCR to test the ability of HERV-K^ΔPBS^-RNAt, HERV-K^ΔPBS^-RNAt-5’LTR, HERV-K^ΔPBS^-RNAt-3’LTR, HERV-K^ΔPBS^-RNAt-5’R, HERV-K^ΔPBS^-RNAt-3’R, HERV-K^ΔPBS^-RNAt-5’R^ΔTBD^ and HERV-K^ΔPBS^-RNAt-3’R^ΔTBD^ RNAs interacting with pTDP-43 protein. Data are shown as mean ± SEM with n=3 biological replicates per experimental condition and a one-way ANOVA with multiple comparisons was performed. *p<0.05; ***p<0.001; ****p<0.0001; ns, no significant difference. **m**, Hypothesized model in which HERV-K RNA triggers nuclear PETANS formation and promotes their release to generate cytosolic TDP-43 aggregates. (made in ©BioRender-biorender.com).

To confirm that these effects were specifically mediated by epigenetically activated endogenous HERV-K, we designed several Dicer-substrate small interfering RNAs (DsiRNAs) targeting endogenous HERV-K transcripts (Fig. 1l). Transfection of these DsiRNAs (DsiRNA1-4) significantly reduced the levels of 5-Aza induced HERV-K transcripts, as validated by RT-qPCR relative to control DsiRNA (Extended Data Fig. 5e). Importantly, co-treatment with these HERV-K targeting DsiRNAs substantially diminished 5-Aza-induced PETANS and cytoplasmic TDP-43 aggregates, whereas control DsiRNA had no such effect (Fig. 1m-o). Collectively, these results indicate that expression of endogenous HERV-K transcripts can promote nuclear PETANS and cytosolic TDP-43 pathology, and that silencing reactivated HERV-K alleviates these phenotypes.

### PETANS are distinct nuclear membraneless organelles

TDP-43 is an RNA and DNA binding protein that contains an intrinsically disordered C-terminal domain and can undergo phase separation in the nucleus in response to some perturbations. For example, expression of an engineered RNA-binding deficient TDP-43 spontaneously demixes to form the outer shell of an intranuclear spherical structure termed the anisosome^38^. The anisosome core is enriched with Hsp70 chaperone family^38^, and its formation depends on Hsp70 activity, as pharmacological inhibitors of Hsp70 disrupts anisosome assembly^39^. Although anisosome formation depends on exogenously expressed TDP-43 that is deficient in RNA binding activity, our findings nevertheless prompted us to ask whether the HERV-K-induced PETANS correspond to the previously described anisosomes. To test this, we first examined if Hsp70 protein was recruited to PETANS as is observed in the case of anisosomes. Immunostaining for endogenous Hsp70 revealed no detectable redistribution or enrichment within PETANS in cells overexpressing HERV-K or HERV-K^ΔPBS^ compared to controls (Fig. 2a). We also examined the localization of an EGFP-Hsp70 fusion protein, previously shown to localize within the core of anisosomes^38^. When we co-expressed this EGFP-Hsp70 fusion along with either HERV-K or HERV-K^ΔPBS^, we found that this ectopic EGFP-Hsp70 was excluded from HERV-K-induced PETANS (Fig. 2b). In contrast to anisosomes, PETANS shows no apparent enrichment for Hsp70.

As a further test of whether PETANS could be the same as anisosomes, we tested the effects of inhibiting Hsp70 function. The Hsp70 inhibitor VER155008 effectively reverses the formation of anisosomes that are induced by overexpressed RNA-binding deficient TDP-43 mutants (TDP-43-5FL or TDP-43-2KQ)^38, 39^. We tested the same treatment to see if it would impact HERV-K-induced PETANS. We first verified that overexpression of the YFP-TDP-43-5FL construct that is deficient in RNA binding could produce anisosomes as previously reported^38, 39^. Indeed, when we expressed this RNA binding deficient TDP-43 for 48 hours in Halo-TDP-43 cells, we detected a robust formation of YFP-tagged anisosomes in the nucleus that also recruited the endogenous Halo-TDP-43 and were positive for pTDP-43 staining (Extended Data Fig. 6). Treatment with 50 µM VER155008 efficiently disrupted these anisosomes, restoring both YFP-TDP-43-5FL and endogenous Halo-TDP-43 to a diffuse nuclear distribution (Extended Data Fig. 6). In contrast, applying the same concentration of VER155008 to HERV-K^ΔPBS^-expressing cells did not disrupt PETANS integrity, and actually led to a slight increase in their nuclear abundance (Fig. 2c-e). These results indicate that Hsp70 exerts distinct effects on the formation or stability of anisosomes versus PETANS, suggesting that PETANS represent a distinct nuclear structure whose formation and contents are mechanistically different from anisosomes, though both are linked to TDP-43 pathology. Unlike anisosomes, PETAN formation takes place with physiological levels of endogenous TDP-43 that has intact RNA binding domains.

We next investigated whether PETANS were associated with any known nuclear membraneless organelles (MLOs). Co-immunostaining using markers of various nuclear MLOs revealed that PETANS partially overlapped with the nucleolus marker, Fibrillarin and with the Cajal body marker, Coilin, though the colocalization was not complete (Extended Data Fig. 7a-c). PETANS did not colocalize with the paraspeckle marker NONO; interestingly, however, NONO was found to colocalized with cytoplasmic TDP-43 aggregates that are labeled by Halo-TDP-43 and pTDP-43 (Extended Data Fig. 7c). These data are consistent with the idea that PETANS are a dynamic structure that can share components with several different nuclear MLOs.

When HERV-K is expressed, nuclear PETANS and cytoplasmic TDP-43 aggregates are observed within the same cells, and in several instances, both structures appeared physically connected across the nuclear envelope (Extended Data Fig. 1e,f). Using real-time live imaging and super-resolution microscopy to track the fluorescent Halo-ligand, we were able to observe nuclear Halo-tagged TDP-43 in HERV-K^ΔPBS^ overexpressing cells. The formation of PETANS appears to be highly dynamic with multiple nascent PETANS precursors emerging as partial spherical assemblies (Fig. 3a, frames 1-15 and Extended Data Movie 1) that later appeared to stabilize (Fig. 3a, frames 20-36). In some instances, nuclear PETANS structures near the nuclear membrane appeared to exit to the cytoplasm (Fig. 3b,c and Extended Data Movies 2,3) to form large TDP-43 inclusions (Fig. 3d and Extended Data Movie 4). In rare circumstances, we also observe PETANS in cytoplasm in fixed cells (Extended data Figure 1e). Further immunostaining analysis revealed that these cytoplasmic TDP-43 aggregates were enriched with nuclear pore complex (NPC) and Lamin B1 proteins (Extended Data Fig. 8a,b), both associated with the nuclear envelope^40^. These findings highlight a possible nuclear origin for these cytosolic aggregates in HERV-K^ΔPBS^-expressing cells, and further support the possibility that nuclear TDP-43 PETANS may contribute to the disruption of nuclear envelope integrity^40^. All of these observations suggest the possibility that cytoplasmic TDP-43 pathology may originate from nuclear PETANS that then translocate into the cytoplasm and act as seeds for cytoplasmic TDP-43 inclusions.

Given that HERV-K expression induced both nuclear PETANS and cytosolic TDP-43 aggregates enriched in pTDP-43, we investigated whether these TDP-43-medated structures also associate with G3BP1 and p62/SQSTM1. G3BP1 is a core stress granule scaffold, and stress granule assembly can facilitate, but is not essential for, cytoplasmic TDP-43 aggregation^41^. In contrast, p62 is a selective autophagy receptor implicated in the recognition and clearance of TDP-43 aggregates^42^. To examine their association with HERV-K-induced TDP-43 structures, we performed immunostaining for G3BP1 and p62 in HERV-K^ΔPBS^-expressing cells and H2B-mCherry-expressing control cells. Neither G3BP1 nor p62 colocalized with nuclear PETANS at any of the time points examined (Extended Data Fig. 9a,b,d,e). By contrast, G3BP1 closely associated with cytosolic TDP-43 aggregates at early time points, but this association progressively declined (Extended Data Fig. 9a,c,g). This decline was accompanied by increasing recruitment of p62 to cytosolic TDP-43 aggregates (Extended Data Fig. 9d,f,g). To further support these findings, we analyzed the soluble and insoluble fractions of TDP-43 in HERV-K expressing cells by Western blotting. Both HERV-K and HERV-K^ΔPBS^ expression promoted a shift in Halo-TDP-43 from the soluble to the insoluble fraction (Extended Data Fig. 9h,i). The insoluble fraction was also enriched in pTDP-43 (Extended Data Fig. 9h,j). Furthermore, HERV-K and HERV-K^ΔPBS^ expression increased cell death in both Halo-TDP-43 and SH-SY5Y cells (Extended Data Fig. 9k,l). Previous studies have reported early colocalization of TDP-43 with stress granules and subsequent recruitment of p62 as cytoplasmic TDP-43 assemblies acquire gel- or solid-like properties^41, 42, 43, 44^. Together, our findings suggest that HERV-K-induced cytosolic TDP-43 aggregates undergo time-dependent molecular remodeling, shifting from dynamic G3BP1-associated assemblies toward p62-associated aggregates, consistent with progressive pathological maturation.

### HERV-K RNA is sufficient to trigger PETANS and cytosolic TDP-43 aggregation

As is the case with XRVs, the HERV-K genomic RNA contains several ORFs encoding structural and enzymatic proteins essential for replication. Expression of HERV-K^ΔPBS^ alone was sufficient to induce nuclear PETANS and cytosolic TDP-43 aggregates across multiple cell types (Fig.1 and Extended Data Fig. 4), suggesting that HERV-K RNA and/or one or more of its coding ORFs triggers these pathological changes in TDP-43. To identify the responsible molecular component, each of the HERV-K ORFs were individually tagged with distinct epitope tags and expressed in Halo-TDP-43 HEK293 cells (Fig. 4a and Extended Data Fig. 10a). None of these ORFs on their own were sufficient to reproduce the effects seen with HERV-K^ΔPBS^ in driving cytoplasmic TDP-43 aggregation (Fig. 4b) or nuclear PETANS formation (Fig. 4c). Although Gag overexpression modestly increased cytoplasmic aggregation above background level (Fig. 4b), it did not significantly induce nuclear PETANS formation (Fig. 4c), implying that Gag might influence TDP-43 pathology through a different mechanism. Collectively, these observations suggest that HERV-K RNA, rather than any of its encoded proteins, triggers both nuclear PETANS and cytosolic TDP-43 aggregates. To further distinguish the contribution of the HERV-K RNA and Gag protein, and to test whether they acted in combination, we generated a HERV-K^ΔPBS^ construct expressing Gag with three tandem Flag tags (HERV-K^ΔPBS^-3FG in Fig. 4d). A second construct, HERV-K^ΔPBS^-3FG-STOP, contained two premature stop codons immediately after the Gag start codon to prevent its translation from HERV-K^ΔPBS^ RNA (Fig. 4d). Immunofluorescent staining confirmed that no detectable Gag expression was found from HERV-K^ΔPBS^-3FG-STOP cells compared with HERV-K^ΔPBS^-3FG (Extended Data Fig. 10b). When these constructs were expressed, they each induced comparable levels of cytosolic Halo-tagged (Fig. 4e,f) or pTDP-43 labelled (Fig. 4e,g) aggregates and showed similar efficiencies in triggering nuclear PETANS (Fig. 4e,h). These data further indicate that HERV-K RNA alone, not the Gag protein, drives PETANS formation and TDP-43 cytoplasmic pathology.

To further delineate which portions of the HERV-K RNA was responsible, we generated a construct in which the entire region that contained each of the coding ORFs was deleted from HERV-K^ΔPBS^, referred to as the HERV-K RNA trigger (HERV-K^ΔPBS^-RNAt in Fig. 5a). Expression of HERV-K^ΔPBS^-RNAt does not contain any HERV-K ORFs and therefore produced only non-coding RNA containing the two LTR domains of HERV-K along with the nuclear mCherry fluorescent protein to visualize successfully transfected cells. When this HERV-K^ΔPBS^-RNAt was expressed in Halo-TDP-43 cells, it induced nuclear PETANS and cytoplasmic TDP-43 aggregation at levels comparable to those observed upon HERV-K^ΔPBS^ expression (Fig. 5b-f). These results further support the idea that HERV-K RNA alone acts as the principal trigger of PETANS and TDP-43 pathology, with the key RNA domain responsible for these phenomena residing within the LTR regions.

We next examined whether HERV-K RNA physically associated with PETANS and cytosolic TDP-43 aggregates. To achieve this, HERV-K^ΔPBS^ was engineered to contain 24 tandem MS2 stem-loop sequences (Extended Data Fig. 11a). Upon expression, the MS2-tagged HERV-K RNA can recruit a GFP tagged MS2 coat protein (MCP) (MCP-GFP), that we co-expressed. This method permits visualization of the subcellular localization of the MS2 stem-loop containing RNA target^45^. Co-expression of MCP-GFP with HERV-K^ΔPBS^-24xMS2 revealed that HERV-K RNA localized to the outer shell of nuclear PETANS and within the cytosolic TDP-43 aggregates, thus co-localizing with both the Halo-tagged TDP43 and pTDP-43 (Extended Data Fig. 11b). This MCP-GFP remained diffusely distributed in control H2B-mCherry-expressing cells (Extended Data Fig. 11b). To further test this physical association between pathological TDP-43 and the HERV-K RNA, we performed RNA immunoprecipitation (RIP) assays using a pTDP-43 antibody to isolate RNA-protein complexes that had been cross-linked by UV irradiation^46^ (Fig. 4i). Total RNA from the pulled-down complexes was analyzed by RT-PCR and RT-qPCR using primer sets targeting the mCherry region that is in common between the control H2B-mCherry, HERV-K-H2B-mCherry and HERV-K^ΔPBS^-H2B-mCherry constructs. 10 ng of a H2B-mCherry DNA construct was used as a normalization control with the same RT-PCR and RT-qPCR conditions. While no H2B-mCherry RNA was detected in pTDP-43 complexes, both HERV-K-H2B-mCherry and HERV-K^ΔPBS^-H2B-mCherry RNAs were strongly enriched (Fig. 4j,k). Notably, HERV-K^ΔPBS^ RNA exhibited stronger interaction with pTDP-43 than thewild-type HERV-K RNA (Fig. 4j,k), consistent with its higher potency in inducing PETANS and cytosolic TDP-43 aggregates (Fig. 1c-g). Together, these findings demonstrate that HERV-K LTR RNA is the principal driver of nuclear PETANS formation and cytosolic TDP-43 aggregates, likely acting through direct physical interactions with pathological TDP-43 assemblies.

We also assessed the critical molecular determinants of *Drosophila* mdg4-ERV in promoting TDP-43 aggregation. Each coding ORF of mdg4-ERV was individually tagged (Extended Data Fig. 12a-c) and co-expressed with human TDP-43-V5. None of these ORFs showed any promotion of cytoplasmic TDP-43 aggregation to a level comparable to that observed with TDP-43 overexpression alone (Extended Data Fig. 12d,e). These results demonstrate that mdg4-ERV ORFs alone are insufficient to promote cytosolic TDP-43 aggregation, highlighting a potential role of mdg4-ERV RNA in driving this process. We next performed RIP to determine whether mdg4-ERV RNA physically associates with pathological TDP-43 complexes in *Drosophila* S2 cells. pTDP-43-containing complexes were immunoprecipitated from cells co-expressing human TDP-43-V5 along with control H2B-mCherry, wild-type mdg4-H2B-mCherry or mdg4^ΔPBS^-H2B-mCherry (Extended Data Fig. 12f). RT-PCR analysis revealed that RNAs from both wild-type mdg4 and mdg4^ΔPBS^ were efficiently pulled down with pTDP-43, whereas no enrichment was observed for the control H2B-mCherry samples (Extended Data Fig. 12g). Notably, sequences corresponding to the LTR and gag regions of mdg4 and mdg4^ΔPBS^ were present in the pTDP-43 complexes, too (Extended Data Fig. 12g). Interestingly, faint LTR and gag signals were also detected in control H2B-mCherry samples. This residual signal likely reflects modest TDP-43 pathology from ectopic human TDP-43-V5 that induced reactivation of endogenous mdg4-ERV from S2 cells (Extended Data Fig. 12g)^9^. Taken together, these findings support the notion that mdg4-ERV RNA, rather than its protein products, acts as the molecular trigger that promotes TDP-43 pathology in a manner analogous to HERV-K RNA. Thus, the ability of ERV RNAs to trigger TDP-43 pathology appears to be an evolutionarily conserved feature of TDP-43’s binding to ERV LTRs.

### The R domain of the HERV-K LTR is sufficient to trigger PETANS and cytosolic TDP-43 pathology

Previous studies have demonstrated that RNA binding promotes TDP-43 liquid-liquid phase separation (LLPS) through sequence-specific interactions, particularly with RNAs containing multiple UG repeat sequences^47^. Sequence analysis revealed that HERV-K RNA contained several UG repeat domains, which could serve as TDP-43 binding sites (Extended Data Fig. 13a). Although several of the HERV-K ORF coding regions contain predicted TDP-43 binding domains (TBDs), expression of these mRNAs failed to recapitulate the ability of full-length HERV-K RNA to induce PETANS and TDP-43 pathology (Fig. 4). So, we focused on the LTR region, comprising the U3, R and U5 elements which are contained within the HERV-K^ΔPBS^-RNAt construct that is sufficient to recapitulate the effects of full length HERV-K. Each of these sub regions of the LTRs harbor several putative TBDs (Extended Data Fig. 13a). To identify the minimal motif within the LTR required for TDP-43 recruitment into PETANS and cytosolic inclusions, we generated a series of truncated LTR constructs (Fig. 5g). Expression of RNAt-5’LTR, RNAt-3’LTR, RNAt-5’R and RNAt-3’R in the Halo-TDP-43 HEK293 cells each elicited PETANS formation and cytosolic TDP-43 aggregates comparable to that of RNAt (Fig. 5h-k). These results indicate that the R region, present in both 5’ and 3’ LTRs, is sufficient to drive these phenotypes, and that R is sufficient to do so whether it is located in the 5’ or 3’ end of a transcript. Sequence inspection revealed a single putative TBD embedded within the R region (Extended Data Fig. 13a). To further test whether this TBD was required for the functional effects on TDP-43, we generated deletion constructs lacking the motif in either the 5’R or 3’R fragment (RNAt-5’R^ΔTBD^ and RNAt-3’R^ΔTBD^ in Fig. 5g). Overexpression of either RNAt-5’R^ΔTBD^ or RNAt-3’R^ΔTBD^ in Halo-TDP-43 cells greatly reduced the activity of the R region to recruit PETANS and induce cytosolic TDP-43 aggregates (Fig. 5h-k). These results demonstrate that the TBD located within the R sub-domain of the LTR of HERV-K RNA is essential for recruiting TDP-43 and promoting both nuclear PETANS and cytosolic TDP-43 aggregates. To further determine whether this identified TBD in R mediated physical interaction between HERV-K RNA and TDP-43, we performed RIP using a pTDP-43 antibody. RT-qPCR analysis of the precipitated complexes showed that deletion of the TBD within R eliminated the interaction between HERV-K RNA and pathological TDP-43 (Fig. 5l). RNA structural analysis of the HERV-K R region also suggested that in principle, R RNAs could be capable of pairing to form inter-molecular complexes (Extended Data Fig. 13b). Multiple sequence alignment of LTRs across HERV families revealed that the TBD involved in TDP-43 binding is specific to HERV-K and absent from other HERV families (Extended Data Fig. 13c,d). Together, these findings indicate a critical TDP-43 binding motif within the R region of HERV-K RNA that mediates its interaction with TDP-43 and that underlies HERV-K-driven nuclear PETANS and cytoplasmic TDP-43 aggregation (Fig. 5m).

## Discussion

One of the hallmarks of ALS, FTD, AD and LATE is the presence of pathological mislocalization of TDP-43 from the nucleus into cytoplasmic inclusions. Such depletion of nuclear TDP-43 results in de-repression of RTEs and ERVs^9, 14, 15, 16, 17, 18, 19^. But such RTE and ERV expression also constitutes a positive feedback to amplify formation of pathological TDP-43 proteinopathy^9^. In this study, we identify a novel pathological process that is triggered in the nucleus by accumulation of HERV-K RNA and that leads to formation of PETANS, a membraneless compartment that contains both pTDP-43 and HERV-K RNA on its outer surface and which is a source of cytosolic TDP-43 inclusions. Thus, the emergence of TDP-43 pathology in the nucleus and then the cytoplasm can be driven by HERV-K RNA expression. Such HERV-K-driven PETANS are observed across multiple human cell lines. It will be important to confirm that such structures are present in human postmortem brain samples, as well as in neurons and glia in animal models. We found that a minimal UG-containing RNA motif within the R region of HERV-K 5’ or 3’ LTR is sufficient to bind TDP-43 and to template both nuclear PETANS formation and cytosolic TDP-43 aggregates. siRNAs that target HERV-K RNA can effectively abolish HERV-K-induced PETANS and cytosolic TDP-43 aggregates. Together, these findings provide mechanistic insight into the previously missing molecular function of dysregulated HERV-K in ALS and FTD patients^17, 20, 21, 22, 23, 24^.

Previous work has shown that total cellular RNA content is critical for regulating phase separation of intrinsically disordered domain containing ribonucleoproteins, including TDP-43. Global nuclear RNA depletion promotes phase separation and suggests that abundant nuclear RNA normally buffers TDP-43 against LLPS and aggregation^11, 48^. In contrast to this system-wide RNA buffering model, other studies have identified distinct TDP-43-binding RNAs that bear multiple TDP-43 binding sites within a single RNA molecule. Such high abundance RNAs can sequester TDP-43 into spatially restricted condensates that provide a locally enriched environment favoring TDP-43 aggregation^49, 50, 51, 52^. We identify HERV-K RNA as being capable of binding and templating TDP-43 aggregates. Mapping of putative TDP-43 binding motifs on HERV-K RNA reveals at least one UG-rich region within the R element of both 5’ and 3’ LTRs. Thus, a single HERV-K RNA transcript presents at least two such R region TDP-43 binding sites and, upon expression, could therefor bring TDP-43 molecules into close proximity to potentiate their aggregation. This concept is in line with prior observations that the ability of RNA to recruit TDP-43 into phase-separated condensates depends on the length of interacting sequences^49, 50^,

^51^. Moreover, RNA structural analysis of the HERV-K R region suggests that R RNA has the potential for self-pairing to form clusters. In principle, this could provide the means to generate higher order RNA assemblies that display multiple TDP-43 binding sites in close proximity and further facilitate TDP-43 aggregate formation (Fig. 5m).

Within the human genome, there are roughly 100 HERV-K proviruses that retain potential coding ORFs and there also are estimated to be around 1000 solo LTR copies^53, 54^. Transcripts derived from either full-length proviruses or solo LTRs contain the TDP-43 binding R element, indicating that a diverse suite of HERV-K loci may contribute to templating TDP-43 pathology at the RNA level. Data from animal models and from postmortem cortical tissues of human subjects indicate that many ERVs and RTEs, including HERV-K, are expressed broadly in neurodegenerative disorders including those that exhibit TDP-43 and Tau pathology^7, 9, 14, 15, 16, 17, 18, 19, 20, 21, 22, 23, 24, 55, 56, 57, 58, 59, 60^. Our previous work in fly models indicates that ERVs contribute functionally to cell intrinsic and intercellular toxic impacts of pathological TDP-43^9, 14, 15^. LINE-1 element expression is also closely associated with both cell intrinsic and intercellular toxicity in mouse models^16^. Based on this growing literature, several ongoing clinical trials are testing antiretroviral therapies, including nucleoside reverse transcriptase inhibitors (NRTIs) to target HERV-K or LINE-1 and integrase inhibitors, to target HERV-K^61, 62^. These strategies rely on the hypothesis that accumulation of cDNA copies of either HERV-K or LINE-1 elements may be a source of inflammatory signaling, for example by activating the cGAS/STING pathway^7, 63, 64^. Our findings do not negate this possibility but provide an additional mechanism by which the RNA transcripts of ERVs may contribute by directly triggering and sustaining the formation of TDP-43 proteinopathy. Our findings therefore underscore the need to develop complementary approaches that directly suppress HERV-K RNA expression, such as the siRNA strategy used in this study, which may improve therapeutic success in TDP-43 associated neurodegenerative diseases.

## Methods

### DNA Constructs

The constructs to express the HERV-K variants were modified from our previously published pcDNA3.1-HERV-K-P2A-H2B-mCherry (abbreviated as HERV-K-H2B-mCherry)^9^. To generate the primer binding site (PBS) deletion construct of HERV-K (HERV-K^ΔPBS^-H2B-mCherry), the following DNA sequence “TGGTGCCCAACGTGGAGG” was deleted from HERV-K-H2B-mCherry construct by PCR. The coding ORFs of HERV-K were PCR-amplified from HERV-K-H2B-mCherry and separately tagged with 3xFlag on Gag (3xFlag-Gag), HA on Pro (Pro-HA), StrepII on Pol (Pol-StrepII) and Myc on Env (Env-Myc). To monitor Gag expression in HERV-K^ΔPBS^-H2B-mCherry, the 3xFlag tag was inserted immediately after the start codon of gag coding sequences to synthesize the HERV-K^ΔPBS^-H2B-mCherry-3FG construct. In order to synthesize no Gag protein expressing construct from HERV-K^ΔPBS^-H2B-mCherry, two repeated stop codon (TAATAG) were placed after the start codon of gag in HERV-K^ΔPBS^-H2B-mCherry-3FG to form HERV-K^ΔPBS^-H2B-mCherry-3FG-STOP. To monitor the cellular localization of HERV-K RNA, 24 tandem copies of MS2 sequences was PCR-amplified from pET264-pUC 24xMS2V6 Loxp KANr Loxp (Addgene #104393) and then integrated into the R-U5 junction of HERV-K^ΔPBS^ to generate HERV-K^ΔPBS^-H2B-mCherry-24xMS2. The MCP-2xGFP that specifically binds to MS2 was PCR-amplified from yWL268 MCP-GFP (Addgene #197052) and then cloned into the pcDNA3.1 vector. To construct RNA trigger (RNAt) of HERV-K (HERV-K^ΔPBS^-H2B-mCherry-RNAt), the entire ORFs sequence was removed from the HERV-K^ΔPBS^-H2B-mCherry. The HERV-K^ΔPBS^-H2B-mCherry-RNAt-5’LTR and HERV-K^ΔPBS^-H2B-mCherry-RNAt-3’LTR were made by separately removing 3’LTR and 5’LTR of HERV-K^ΔPBS^-H2B-mCherry-RNAt. To construct HERV-K^ΔPBS^-H2B-mCherry-RNAt-5’R and HERV-K^ΔPBS^-H2B-mCherry-RNAt-3’R, the sequences correlated with U5 in HERV-K^ΔPBS^-H2B-mCherry-RNAt-5’LTR, and U3 and U5 in HERV-K^ΔPBS^-H2B-mCherry-RNAt-3’LTR were deleted by PCR. The potential TDP-43 binding domain (TGTTTATGTGTATG) in R region of HERV-K was separately deleted in HERV-K^ΔPBS^-H2B-mCherry-RNAt-5’R and HERV-K^ΔPBS^-H2B-mCherry-RNAt-3’R constructs to generate the HERV-K^ΔPBS^-H2B-mCherry-RNAt-5’R^ΔTBD^ and HERV-K^ΔPBS^-H2B-mCherry-RNAt-3’R^ΔTBD^. The constructs expressing EGFP-Hsp70 (Addgene #15215) was obtained from the Addgene.

The pAc5.1-H2B-mCherry, pMT-TDP-43-V5 and mdg4 expressing constructs were published or modified from our previously published pAc5.1-mdg4-P2A-H2B-mCherry (abbreviated as mdg4-H2B-mCherry)^9, 65^. The PBS sequence (TGGCGCCCAAC) of mdg4 was deleted from mdg4-H2B-mCherry to synthesize the mdg4^ΔPBS^-H2B-mCherry. The coding ORFs of mdg4 were PCR-amplified from mdg4-P2A-H2B-mCherry and separately tagged with Flag on Gag (Gag-Flag), StrepII on Pol (Pol-StrepII) and Myc on Env (Env-Myc).

### Cell lines and cell culture

The CRISPR-engineered HaloTag-TDP-43 (Halo-TDP-43)^25^, HaloTag-GAPDH (Halo-GAPDH)^25^ and wild-type HEK293T (ATCC, CRL-11268) cells were cultured in Dulbecco’s Modified Eagle Medium (DMEM) (Thermo Fisher Scientific, 11995065). Human neuroblastoma cells, SH-SY5Y (ATCC, CRL-2266) was maintained in DMEM/Nutrient Mixture F-12 (DMEM/F-12) (Thermo Fisher Scientific, 11320033), and human microglia cells HMC3 (ATCC, CRL-3304) and human astrocytes SVG p12 (ATCC, CRL-8621) were cultured in Eagle’s Minimum Essential Medium (EMEM) in a 37°C incubator with 5% CO_2_ supplementation. All mammalian cell media were supplemented with 10% fetal bovine serum (FBS) (Thermo Fisher Scientific, 10438026) and 100 U/ml Penicillin-Streptomycin (Thermo Fisher Scientific, 15140122). *Drosophila* S2 cells (Thermo Fisher Scientific, R69007) were grown in Schneider’s *Drosophila* medium (Thermo Fisher Scientific, 21720001) supplemented with 10% FBS and 100U/ml Penicillin-Streptomycin at 25°C.

### Transfection of culture cells

A total of 5×10^5^ Halo-TDP-43 or wild-type HEK293T cells were seeded onto Poly-D-Lysine-coated coverslips (Electron Microscopy Sciences, 72294-16) placed in 6-well culture plates and incubated overnight before transfection. Transfection was carried out according to the manufacturer’s instructions using a mixture of 4 µg plasmid DNA and 12 µl TransIT-2020 reagent (Mirus, MIR 5400) at a 1:3 ratio. Twenty-four hours post-transfection, the supernatant containing residual transfection reagent was removed, and the cells were washed twice with 1xPBS. Fresh complete medium was replenished daily until the designated time point for analysis. SH-SY5Y, HMC3 and SVG p12 cells were seeded at a density of 1×10^5^ cells per well onto coverslips coated with 0.1 mg/ml Poly-D-Lysin (Thermo Fisher Scientific, A3890401). Transfection conditions and subsequent analyses were performed as described above for Halo-TDP-43.

Dicer-Substrate siRNAs (DsiRNA) were synthesized by Integrated DNA Technologies (IDT). Each DsiRNA was transfected at a final concentration of 10 nM using TransIT-2020 reagent (Mirus, MIR 5400) at a 1:3 ratio. The sequences of each DsiRNA were listed as follows: DsiRNA-1 (sense 5’-rGrCrUrUrUrGrUrArArArGrCrArUrUrGrArGrArUrGrUrUTA-3’ and antisense 5’-rUrArArArCrArUrCrUrCrArArUrGrCrUrUrUrArCrArArArGrCrArG-3’), DsiRNA-2 (sense 5’-rArCrArUrUrGrUrCrUrArUrGrArUrGrCrArArArGrArCrCTT-3’ and antisense 5’-rArArGrGrUrCrUrUrUrGrCrArUrCrArUrArGrArCrArArUrGrUrArA-3’), DsiRNA-3 (sense 5’-rGrArGrArUrUrCrUrGrUrUrArArUrCrUrArUrGrArCrCrUTA-3’ and antisense 5’-rUrArArGrGrUrCrArUrArGrArUrUrArArCrArGrArArUrCrUrCrArA-3’), DsiRNA-4 (sense 5’-rGrGrUrUrUrCrUrCrCrCrCrArUrGrUrGrArUrArGrUrCrUGA-3’ and antisense 5’-rUrCrArGrArCrUrArUrCrArCrArUrGrGrGrGrArGrArArArCrCrUrU-3’) and control DsiRNA (IDT, 51-01-19-09).

S2 cells transfection was carried out following the conditions described in our previous study^9^. Briefly, 5×10^5^ S2 cells were seeded onto coverslips coated with 0.5 mg/ml concanavalin A (ConA) (MP Biomedicals, 195283) and placed in 6-well culture plates overnight prior to transfection. Co-transfections with the indicated constructs (pAc5.1-H2B-mCherry, pAc5.1-mdg4-H2B-mCherry, pAc5.1-mdg4^ΔPBS^-H2B-mCherry and pMT-hTDP-43-V5) were performed using 1 µg of each plasmid DNA and Effectene transfection reagent (Qiagen, 301427). Twenty-four hours after transfection, transfection complexes were washed away, and transfected cells were supplied with the fresh complete medium containing 700 µM CuSO_4_ to induce hTDP-43-V5 expression.

### Monitoring of Halo-tagged proteins

Transfected cells were incubated with Janelia Fluor 646 HaloTag Ligand (Promega, GA1120) at final concentration of 200 nM prior to final analysis. Cells incubated with HaloTag labeling were performed at 37°C for 30 minutes. To minimize background fluorescence from unbound ligand, the medium was replaced with fresh complete medium and cells were incubated at 37°C for 15 minutes. This washing step was repeated twice. Following HaloTag labeling, the cells were subjected to subsequent analyses.

### Drug treatment

5-Aza (Millipore-Sigma, A1287) was prepared at final concentrations of 500 nM and 1000 nM. During transfection, cells were simultaneously treated with 5-Aza. The medium containing 5-Aza was refreshed daily, and cells were harvested at the indicated time points for subsequent analyses. VER155008 (Bio-Techne, 3803) was prepared at a final concentration of 50 µM. One day after transfection, the culture medium containing with transfection reagents was replaced with medium supplemented with VER155008, which was refreshed daily until desired time point for analysis.

### Immunostaining of transfected cells

Mammalian cells or *Drosophila* S2 cells grown on Poly-D-Lysine or ConA coated coverslips were washed three times with 1xPBS and immediately fixed in 4% paraformaldehyde (Electron Microscopy Sciences #15713) PBS solution (1xPBS with 4% paraformaldehyde and 0.2% Triton-X-100 (SIGMA-ALDRICH)) for 10 minutes on a nutator. After fixation, cells were washed three times with 1xPBST wash solution (1xPBS with 1% Triton-X-100 and 3% NaCl) and then incubated for 30 minutes in blocking solution (1xPBST with 10% normal horse serum) at room temperature. Following blocking, cells were incubated for 1 hour at room temperature with primary antibodies diluted in 1xPBST washing solution containing 10% normal horse serum: rabbit anti-pTDP-43 (1:350, proteintech 22309-1-AP), mouse anti-TDP-43 (1:100, proteintech 60019-2-Ig), mouse anti-Hsp70 (1:100, abcam ab2787), mouse anti-Fibrillarin (1:100, Thermo Fisher Scientific MA3-16771), mouse anti-NONO (1:100, Thermo Fisher Scientific MA3-2024), mouse anti-Coilin (1:100, abcam ab87913), mouse anti-FUS (1:100, proteintech 68262-1-Ig), mouse anti-hnRNP A1 (1:100, Santa Cruz sc-32301), mouse anti-NPC (1:200, abcam ab24609), mouse anti-Lamin B1 (1:200, proteintech 66095-1-Ig), mouse anti-G3BP1 (1:100, abcam ab56574), mouse anti-p62 (1:100, Cell Signaling Technology 88588S), rabbit anti-p62 (1:100, proteintech 18420-1-AP), mouse anti-Flag (1:250, Millipore-Sigma F3165), rat anti-HA (1:200, Roche 11867423001), rat anti-StrepII (1:100, abcam ab252885), rat anti-Myc (1:200, abcam ab10910) and mouse anti-V5 (1:250, Thermo Fisher Scientific R960-25). After primary antibody incubation, samples were washed three times with 1xPBST for 10 minutes each and then incubated with secondary antibodies solution (1xPBST washing solution with 10% normal horse serum) with DAPI at room temperature for 1 hour. Cells were washed again three times with 1xPBST for 10 minutes before mounting with ProLong Diamond Antifade Mountant (Thermo Fisher Scientific, P36961). Imaging was performed using a Zeiss LSM 800 confocal microscope with Airyscan mode, and images were processed using Zeiss Zen software package.

### Live cell imaging

Cells were seeded onto Poly-D-Lysine-coated glass bottom 35 mm dish (Thermo Fisher Scientific, 150680) and transfected with HERV-K^ΔPBS^-H2B-mCherry under the conditions described above. At the designated time point, cells were incubated with HaloTag Ligand. Following incubation, the HaloTag-Ligand containing medium was removed, and the cells were washed and replenished with fresh FluoroBrite DMEM supplemented with 10% FBS and 100 U/ml Penicillin-Streptomycin. Live-cell imaging was immediately performed using a Zeiss LSM 980 confocal microscope equipped with Airyscan 2 and two-photon scanning capabilities.

### Image analysis

To quantify cell populations exhibiting nuclear PETANS, cytoplasmic Halo or pTDP-43 aggregates, images covering an area of 25513.68 µm^2^ (159.73 µm x 159.73 µm) were collected. The percentage of cells with nuclear PETANS was calculated as the number of PETANS^+^ and H2B-mCherry^+^ double-positive cells divided by the total number of H2B-mCherry^+^ cells within the imaged area. The same criterion was applied to determine the proportion of cells with cytoplasmic Halo or pTDP-43 aggregates. In 5-Aza-treated samples, PETANS^+^, cytoplasmic Halo^+^ and pTDP-43^+^ cells were quantified and normalized to the total number of DAPI-positive cells to evaluate the effect of 5-Aza treatment on these phenomena. To quantify the total number of PETANS within individual nuclei, super-resolution images of single H2B-mCherry^+^ cells were acquired, and PETANS counts were determined from these images. The three-dimensional displays of the images were generated using Imaris software (Oxford Instruments).

For quantification of co-localization with FUS, hnRNP A1, NPC, Lamin B1, G3BP1 and p62, cells were classified as positive if at least one cytoplasmic TDP-43 aggregate displayed co-localization with the indicated protein, regardless of whether all cytosolic aggregates within that cell were double-positive.

### RNA immunoprecipitation (RIP)

Cells were washed once with ice-cold 1xPBS, followed by the addition of 2 mL ice-cold 1xPBS to each well of a 6-wull culture plate. The plate lid was removed and placed on the ice. Cells were UV-crosslinked twice using a UV crosslinker (Stratagene Model 1800) at 250 mJ. The cells were then collected into 15 mL centrifuge tubes and centrifuged at 300xg for 5 minutes. After removing the supernatant, the cell pellets were placed on dry ice for 10 minutes and stored at −80°C until further use. For lysis, cell pellets were resuspended in 1 mL lysis buffer (50 mM Tris-HCl, pH 8.0; 100 mM NaCl; 1% NP-40; protease inhibitor cocktail (Roche, 11873580001), phosphatase inhibitor cocktail (Roche, 4906845001) and 50 U/mL RNase inhibitor (Promega, N2611)) and incubated on ice for 30 minutes. The cell lysates were centrifuged at 13000xg for 20 minutes at 4°C, and the supernatant was transferred to Dynabeads preincubated with 2 µg of anti-pTDP-43 antibody. Samples were incubated with rotation at 4°C for 2 hours. After incubation, the supernatant was discarded, and the beads were washed twice with wash buffer (20 mM Tris-HCl, pH 7.4; 10 mM MgCl_2_; 0.5% NP-40; protease and phosphatase inhibitors and 50 U/mL RNase inhibitor). RNA was isolated from the beads using TRIzol reagent (Thermo Fisher Scientific, 15596026) protocol. The isolated RNA was either used for immediately for downstream analyses or stored at −80°C until use.

### RT-qPCR for HERV-K, TDP-43 pathology target and RIP complexes

Total RNA was extracted from Halo-TDP-43 cells using TRIzol reagent (Thermo Fisher Scientific, 15596026) following the manufacturer’s protocol as previously described. The isolated RNA was subsequently treated with RQ1 DNase (Promega, M6101) to remove residual DNA prior to reverse transcription. Complementary DNA was synthesized from 0.5 µg of total RNA using the PrimeScript RT Reagent kit (TaKaRa, RR037A). qPCR was carried out with PowerUp SYBR Green Master Mix (Thermo Fisher Scientific, A25742) on a QuantStudio 3 real-time PCR system. The primer sets used for target amplification were as follows: HERV-K LTR5_Hs (forward 5’-GGGCAGCAATACTGCTTTGT-3’ and reverse 5’-CAATAGTGGGGAGAGGGTCA-3’), HERV-K gag (forward 5’-AAATAAGACCCAACCGCCAGTAGC-3’ and reverse 5’-GAATTGCCATGCCTCAGTATCTCC-3’), UNC13A-CE (forward 5’-TGGATGGAGAGATGGAACCT-3’ and reverse 5’-GGGCTGTCTCATCGTAGTAAAC-3’)^30, 31^, STMN2-CE (forward 5’-GCTCTCTCCGCTGCTGTAG-3’ and reverse 5’-CTGTCTCTCTCTCTCGCACA-3’)^30^, GAPDH (forward 5’-GTTCGACAGTCAGCCGCATC-3’ and reverse 5’-GGAATTTGCCATGGGTGGA-3’), mCherry 1 (forward 5’-CAGAGGCTGAAGCTGAAGGA-3’ and reverse 5’-CACCTACAAGGCCAAGAAGC-3’), mCherry 2 (forward 5’-GCGTGATGAACTTCGAGGAC-3’ and reverse 5’-AGAAGAAGACCATGGGCTGG-3’), mdg4 LTR (forward 5’-ACAACTAACAATGTATTGCTTCGTAG-3’ and reverse 5’-TTTCGCTGAGGTTCGTCTTAG-3’), and mdg4 gag (forward 5’-AACTCAATTCCCTAGCTGCG-3’ and reverse 5’-TCAGCACCCTCTTGTTCATG-3’).

### Western Blot analysis for soluble and insoluble fractions of TDP-43

The soluble and insoluble TDP-43 fractions from Halo-TDP-43 cells were sequentially extracted dependent on the solubilities in the extraction buffers as previously described^9, 42^. Transfected Halo-TDP-43 cells were washed, collected with 1XPBS, and then homogenized in ice-cold RIPA buffer (Thermo Fisher Scientific, 89900) with protease and phosphatase inhibitors (Thermo Fisher Scientific, A32961). Cell lysates were centrifuged at 16000g for 30 minutes at 4°C, and the supernatant was saved as the RIPA-soluble fraction. The RIPA-insoluble pellets were washed with RIPA buffer by centrifuging at 16000g for 15 minutes at 4°C. Supernatants were discarded and the pellets were extracted and homogenized with urea buffer (7M Urea, 4% CHAPS, 2M Thiourea, 30mM Tris, pH=8.5). The lysates were centrifuged at 16000g for 30 minutes at 4°C. The supernatant was kept as the RIPA-insoluble and urea-soluble fraction.

30µg protein samples from each cellular fraction were separated by 10% polyacrylamide gel (BIO-RAD, 4561034) and followed by a canonical Western Blot protocol. In short, membranes were blocked with 5% bovine serum albumin (BSA) (GoldBio, A-420-100) in 1XTBS (BIO-RAD, 1706435) with 0.2% of Tween-20 (TBST) at room temperature for 30 minutes. After blocking, membranes were incubated with primary antibodies mouse anti-TDP-43 (1:1000, proteintech 60019-2-Ig), rabbit anti-pTDP-43 (1:1000, proteintech 22309-1-AP), mouse anti-GAPDH (1:5000, Santa Cruz Biotechnology sc-47724), and rabbit anti-αTubulin (1:5000, Cell Signaling Technology 9099S) in 5% BSA-TBST for 1 hour at room temperature or overnight at 4°C. After washing three times with 1XTBST, membranes were incubated with secondary antibodies, donkey anti-mouse IRDye 800CW (LICORbio, 926-32212) or donkey anti-rabbit IRDye 680RD (LICORbio, 926-68073) at 1:2000 or 1:5000 dilutions), for 1 hour at room temperature. Signals were imaged with Sapphire Biomolecular Imager (Azure Biosystems).

### Cell viability and death assays

Transfected cells were trypsinized at identical time points, resuspended in culture medium, and mixed with trypan blue (Thermo Fisher Scientific, 15250061). Viable and dead cells were then counted using a hemocytometer under microscopy. Cell viability was also assessed using the CellTiter-Glo 2.0 assay (Promega, G9242) following the manufacturer’s protocol. Briefly, CellTiter-Glo 2.0 reagent was added to each well at 1:1 ratio with the culture medium. After shaking for 2 minutes to promote cell lysis, plates were incubated at room temperature for 10 minutes to stabilize the luminescent signal. Luminescence was then quantified using a luminometer (Molecular Devices Filter Max F5).

### Multiple sequence alignment of TBD motifs across HERV RNAs

To assess the conservation of the TBD across HERV families, LTR consensus sequences were obtained from Repbase (Version 2.23) and subjected to multiple sequence alignment (MSA) using EMBL-EBI Clustal Omega. The initial analysis included LTR1 to LTR20 sequences. Sequences lacking regions alignable to the TBD domain were excluded from subsequent analysis. Eight families, including HERV-K, with relatively conserved sequences were retained for a second round of alignment.

### Statistics and reproducibility

Statistical analyses were performed using GraphPad Prism 10. Details of the statistical tests, significance comparisons between experimental groups, and measures of experimental reproducibility were provided in the corresponding figure legends. Significant levels were denoted as follows: *p<0.05, **p<0.01, ****p<0.001 and ****p<0.0001.

## Data availability

All data are available in the main text and the supplementary materials. All newly created reagents are available upon request with a Materials transfer agreements (MTAs).

## Code availability

This study did not generate/analyze code.

## Acknowledgment

We thank Professor Sami Barmada (University of Michigan) for providing us the Halo-TDP-43 and Halo-GAPDH cell lines. We thank all members of the Sher, Nelson and Dubnau labs for helpful discussions. This work was supported by NIA grants R01AG078788 and R01AG076493 to J.D. We also are grateful to Andrew Frank, Roger Sher and Anthony Zador and for critical comments on the manuscript.

## Author contributions

Y.H.C. and J.D. designed the experiments. Y.H.C. performed all experiments, collected and analyzed the data. Y.H.C. and J.D. wrote the manuscript.

## Figure legends

**Extended Data Fig. 1 HERV-K and HERV-K expression and validation. a**, Schematic illustration of primer sets used to detect HERV-K expression. RT-qPCR analysis was performed to validate the expression of HERV-K and HERV-K^ΔPBS^ using primer sets located in the regions of **b**, LTR and **c**, gag. Data are shown as mean ± SEM with n=3 biological replicates per timepoint and a two-way ANOVA with multiple comparisons was performed. *p<0.05; **p<0.01; ***p<0.001; ****p<0.0001; ns, no significant difference. **d**, RT-qPCR analysis was performed to validate the expression of endogenous TDP-43 after HERV-K and HERV-K^ΔPBS^ overexpression. Data are shown as mean ± SEM with n=3 biological replicates per timepoint and a two-way ANOVA with multiple comparisons was performed. ns, no significant difference. **e**, PETANS were occasionally detected in the cytosol (pink arrowhead). Halo-TDP-43 (cyan), pTDP-43 (green), H2B-mCherry (red), and DAPI (blue). f, view of PETANS in X, Y and Z planes to provide three-dimensional view. The three-dimensional view of PETANS and cytosolic TDP-43 aggregate. **g**, Average diameter of PETANS from super-resolution images of Halo-TDP-43 cells expressing HERV-K (0.99±0.09 µm, n=19 PETANS) or HERV-K^ΔPBS^ (1.19±0.08 µm, n=58 PETANS). Data are shown as mean ± SEM.

**Extended Data Fig. 2 Co-localization of FUS and hnRNP A1 with nuclear PETANS and cytosolic TDP-43 aggregates following HERV-K**^**ΔPBS**^ **expression. a**, Subcellular distribution of FUS with nuclear PETANS and cytosolic TDP-43 aggregates were visualized by Halo-TDP-43 (cyan), pTDP-43 (blue), H2B-mCherry (red), and FUS (green) in response to control H2B-mCherry or HERV-K^ΔPBS^ expression at 72 hours. Scale bar=5 µm. Quantification of FUS co-localization in cells exhibiting nuclear PETANS and cytosolic TDP-43 aggregates. FUS co-localization was detected in cells with PETANS, 0.00±0.00 (n=58) and in cells with TDP-43 aggregates, 0.23±0.06 (n=47). **b**, Subcellular distribution of hnRNP A1 with nuclear PETANS and cytosolic TDP-43 aggregates were visualized by Halo-TDP-43 (cyan), pTDP-43 (blue), H2B-mCherry (red), and hnRNP A1 (green) after control H2B-mCherry or HERV-K^ΔPBS^ expression for 72 hours. Scale bar=5 µm. Quantification of hnRNP A1 co-localization in cells exhibiting nuclear PETANS and cytosolic TDP-43 aggregates. hnRNP A1 co-localization was detected in cells with PETANS, 0.00±0.00 (n=48) and in cells with TDP-43 aggregates, 0.05±0.04 (n=37).

**Extended Data Fig. 3 Co-expression *Drosophila* mdg4 and mdg4**^**ΔPBS**^ **with human TDP-43. a**, Schematic description of control H2B-mCherry, mdg4 (mdg4-H2B-mCherry) and mdg4^ΔPBS^ (mdg4^ΔPBS^-H2B-mCherry) constructs. **b**, Illustration of the timeframe for co-transfection and imaging. c, Experimental images of co-transfection of control H2B-mCherry, mdg4 and mdg4^ΔPBS^ with human TDP-43-V5 with staining for V5 (green), pTDP-43 (yellow), H2B-mCherry (red) and DAPI (blue). Scale bar=5 µm. Quantification of population percentage of cells showing cytoplasmic **d**, pTDP-43 (H2B-mCherry 19.44±2.41%, n=31; mdg4 50.07±2.93%, n=30 and mdg4^ΔPBS^ 43.15±3.51%, n=21) and **e**, TDP-43-V5 (H2B-mCherry 32.34±3.50%, n=31; mdg4 69.71±4.12%, n=30 and mdg4^ΔPBS^ 59.98±4.53%, n=21) aggregation. Data are shown as mean ± SEM and a one-way ANOVA with multiple comparisons was performed. ****p<0.0001; ns, no significant difference.

**Extended Data Fig. 4 HERV-K expression causes PETANS and cytosolic TDP-43 aggregates in Halo-GAPDH, wild-type HEK293T, microglia, astrocyte and neuroblastoma cells. a**, Immunofluorescent images of Halo-GAPDH HEK293T cells expressing control H2B-mCherry and HERV-K with staining for Halo-GAPDH (cyan), pTDP-43 (green), H2B-mCherry (red) and DAPI (blue). Scale bar=5 µm. **b**, Images of wild-type HEK293T cells expressing control H2B-mCherry, HERV-K and HERV-K^ΔPBS^ with staining for pan-TDP-43 (cyan), pTDP-43 (green), H2B-mCherry (red) and DAPI (blue). Scale bar=5 µm. **c**, Images of HMC3 microglial cells expressing control H2B-mCherry, HERV-K and HERV-K^ΔPBS^ with staining for pan-TDP-43 (cyan), pTDP-43 (green), H2B-mCherry (red) and DAPI (blue). Pink arrowhead indicates PETANS and yellow arrowhead indicates cytosolic TDP-43 aggregate. Scale bar=5 µm. **d**, Images of SVG p12 astrocyte expressing control H2B-mCherry, HERV-K and HERV-K^ΔPBS^ with staining for pan-TDP-43 (cyan), pTDP-43 (green), H2B-mCherry (red) and DAPI (blue). Pink arrowhead indicates PETANS and yellow arrowhead indicates cytosolic TDP-43 aggregate. Scale bar=5 µm. **e**, Images of SH-SY5Y neuroblastoma expressing control H2B-mCherry and HERV-K with staining for pan-TDP-43 (cyan), pTDP-43 (green), H2B-mCherry (red) and DAPI (blue). Pink arrowhead indicates PETANS and yellow arrowhead indicates cytosolic TDP-43 aggregate. Scale bar=5 µm.

**Extended Data Fig. 5 Reactivation of endogenous HERV-K by 5-Aza. a**, Schematic illustration of the timeframe and effect of 5-Aza treatment on Halo-TDP-43 cells. **b**, Validation of 5-Aza-mediated reactivation of HERV-K by RT-qPCR. Data are shown as mean ± SEM with n=3 per timepoint and a one-way ANOVA with multiple comparisons was performed. *p<0.05; ****p<0.0001; ns, no significant difference. **c**, Images of 5-Aza-triggered PETANS and cytosolic TDP-43 aggregate in Halo-TDP-43 cells with staining for pan-TDP-43 (cyan), pTDP-43 (green) and DAPI (blue). Scale bar=5 µm. **d**, 5-Aza-induced CE inclusion in *STMN2* and *UNC13A* was assessed by RT-qPCR. Data are shown as mean ± SEM with n=3 per timepoint. A one-way ANOVA with multiple comparisons was performed. **p<0.01; ns, no significant difference. **e**, Validation of the effects of DsiRNAs targeting HERV-K on 1000 nM 5-Aza-mediated reactivation of HERV-K by RT-qPCR on LTR5_Hs and gag levels. Data are shown as mean ± SEM with n=3 per timepoint and a one-way ANOVA with multiple comparisons was performed. **p<0.01; ***p<0.001; ****p<0.0001; ns, no significant difference.

**Extended Data Fig. 6 Hsp70 inhibitor dissolved TDP-43-5FL-induced anisosomes. a**, Schematic of the timeframe of Hsp70 inhibitor treatment on YFP-TDP-43-5FL induced anisosomes. **b**, Immunofluorescent images showing the effect of the Hsp70 inhibitor on YFP-TDP-43-5FL-induced anisosomes, staining for Halo-TDP-43 (cyan), pTDP-43 (white), YFP-TDP-43-5FL (yellow), and H2B-mCherry (red). Scale bar=5 µm. **c**, Cell population with YFP-TDP-43-5FL-induced anisosomes with Hsp70 inhibitor treatment. (DMSO 71.85±5.00%, n=30 and VER155008 3.54±1.68%, n=36). Data are shown as mean ± SEM and unpaired t-test was performed. ****p<0.0001. **d**, Difference in YFP-TDP-43-5FL-induced anisosome number after Hsp70 inhibitor treatment from super-resolution 3D images. (DMSO 20.29±3.33%, n=7 and VER155008 1.08±0.73%, n=12). Data are shown as mean ± SEM and unpaired t-test was performed. ****p<0.0001.

**Extended Data Fig. 7 Co-staining of several nuclear MLO markers with PETANS. a**, Co-staining of the nucleolus marker Fibrillarin with HERV-K-induced PETANS and the colocalization ratio of Fibrillarin and PETANS (HERV-K 29.49±5.20%, n=78 PETANS and HERV-K^ΔPBS^ 19.05±6.13%, n=42 PETANS). Halo-TDP-43 (cyan), pTDP-43 (blue), Fibrillarin (green), and H2B-mCherry (red). Data are shown as mean ± SEM. Scale bar=5 µm. **b**, Co-staining of the Cajal body marker Coilin with HERV-K-induced PETANS and the colocalization ratio of Coilin and PETANS (HERV-K 31.37±6.56%, n=51 PETANS and HERV-K^ΔPBS^ 31.67±6.06%, n=60 PETANS). Halo-TDP-43 (cyan), pTDP-43 (blue), Coilin (green), and H2B-mCherry (red). Data are shown as mean ± SEM. Scale bar=5 µm. c, Co-staining of the paraspeckle marker NONO with HERV-K-induced PETANS and the colocalization ratio of NONO and PETANS (HERV-K 15.38±5.85%, n=39 PETANS and HERV-K^ΔPBS^ 10.91±4.24%, n=55 PETANS). Halo-TDP-43 (cyan), pTDP-43 (blue), NONO (green), and H2B-mCherry (red). Data are shown as mean ± SEM. Scale bar=5 µm.

**Extended Data Fig. 8 Co-localization of NPC and Lamin B1 with nuclear PETANS and cytosolic TDP-43 aggregates following HERV-K**^**ΔPBS**^ **expression. a**, Subcellular distribution of NPC with cytosolic TDP-43 aggregates were visualized by Halo-TDP-43 (cyan), pTDP-43 (blue), H2B-mCherry (red), and NPC (green) after control H2B-mCherry or HERV-K^ΔPBS^ expression for 72 hours. Scale bar=5 µm. Quantification of NPC co-localization in cells exhibiting cytosolic TDP-43 aggregates. NPC co-localization was detected in cells with TDP-43 aggregates, 0.71±0.06 (n=59). **b**, Subcellular distribution of Lamin B1 with cytosolic TDP-43 aggregates were visualized by Halo-TDP-43 (cyan), pTDP-43 (blue), H2B-mCherry (red), and Lamin B1 (green) after control H2B-mCherry and HERV-K^ΔPBS^ expression for 72 hours. Scale bar=5 µm. Quantification of Lamin B1 co-localization in cells exhibiting cytosolic TDP-43 aggregates. Lamin B1 co-localization was detected in cells with TDP-43 aggregates, 0.59±0.09 (n=34).

**Extended Data Fig. 9 Co-localization of G3BP1 and p62 with nuclear PETANS and cytosolic TDP-43 aggregates following HERV-K**^**ΔPBS**^ **expression. a**, Subcellular distribution of G3BP1 with nuclear PETANS and cytosolic TDP-43 aggregates were visualized by Halo-TDP-43 (cyan), pTDP-43 (white), H2B-mCherry (red), and G3BP1 (green) after control H2B-mCherry or HERV-K^ΔPBS^ expression for 72 or 120 hours. Scale bar=5 µm. Quantification of G3BP1 co-localization in cells exhibiting **b**, nuclear PETANS and **c**, cytosolic TDP-43 aggregates. G3BP1 co-localization was detected in cells with PETANS, (0.00±0.00 (n=69) at 72hrs and 0.00±0.00 (n=54) at 120hrs) and in cells with TDP-43 aggregates, (0.79±0.05 (n=66) at 72hrs and 0.74±0.06 (n=53) at 120hrs). **d**, Subcellular distribution of p62 with nuclear PETANS and cytosolic TDP-43 aggregates were visualized by Halo-TDP-43 (cyan), pTDP-43 (white), H2B-mCherry (red), and p62 (green) after control H2B-mCherry or HERV-K^ΔPBS^ expression for 72 vs 120 hours. Scale bar=5 µm. Quantification of p62 co-localization in cells exhibiting **e**, nuclear PETANS and **f**, cytosolic TDP-43 aggregates. p62 co-localization was detected in cells with PETANS, (0.00±0.00 (n=38) at 72hrs and 0.00±0.00 (n=25) at 120hrs) and in cells with TDP-43 aggregates, (0.27±0.08 (n=36) at 72hrs and 0.57±0.09 (n=33) at 120hrs). **g**, Subcellular distribution of G3BP1 and p62 with nuclear PETANS and cytosolic TDP-43 aggregates were visualized by Halo-TDP-43 (cyan), G3BP1 (white), H2B-mCherry (red), and p62 (green) under control H2B-mCherry and HERV-K^ΔPBS^ expression at 120 hours. Scale bar=5 µm. **h**, The distribution of Halo-TDP-43 and pHalo-TDP-43 in soluble (RIPA; R) and insoluble (Urea; U) fractions under control H2B-mCherry, HERV-K and HERV-K^ΔPBS^ expression was analyzed by Western Blot. GAPDH and α-Tubulin were used as markers for soluble fraction. Quantification of **i**, the ratio of insoluble (U) to soluble (R) Halo-TDP-43 and **j**, fold change of pHalo-TDP-43 derived from the Western Blot analysis in panel h. Cell viability and cell death were assessed using trypan blue exclusion assay and CellTiter-Glo 2.0 luminescence viability measurements of **k**, Halo-TDP-43 and l, SH-SY5Y cells under control H2B-mCherry, HERV-K and HERV-K^ΔPBS^ expression.

**Extended Data Fig. 10 Characterization of HERV-K ORFs expression and Gag expression in HERV-K**^**ΔPBS**^ **-3FG and HERV-K**^**ΔPBS**^ **-3FG-STOP constructs. a**, Validation of the expression constructs for HERV-K ORFs with different tags fused with Gag, Pro, Pol, and Env, generating 3xFlag-Gag (yellow), Pro-HA (yellow), Pol-StrepII (yellow), and Env-Myc (yellow) for visualization in expressing cells, co-stained with Halo-TDP-43 (cyan), H2B-mCherry (red), and DAPI (blue). Scale bar=5 µm. **b**, Characterization of Gag expression from control H2B-mCherry, HERV-K^ΔPBS^-3FG and HERV-K^ΔPBS^-3FG-STOP constructs in Halo-TDP-43 cells, co-stained for Flag (green), Halo-TDP-43 (cyan), pTDP-43 (white), and H2B-mCherry (red). Scale bar=5 µm.

**Extended Data Fig. 11 Visualization of HERV-K RNA by MS2-MCP system. a**, Construct design of HERV-K^ΔPBS^-MS2. **b**, Visualization of the subcellular localization of HERV-K RNA in Halo-TDP-43 cells with control H2B-mCherry and HERV-K^ΔPBS^-MS2 co-transfected with MCP-GFP, staining with Halo-TDP-43 (cyan), pTDP-43 (yellow), MCP-GFP (green), and H2B-mCherry (red). Scale bar=5 µm.

**Extended Data Fig. 12 Characterization of mdg4 ORF and RNA expression and their effects on TDP-43 aggregate in *Drosophila* S2 cell. a**, Construct design to visualize mdg4 ORFs. **b**, Schematic illustration of the experimental timeframe testing the effect of mdg4 ORFs on TDP-43 aggregation. **c**, Validation the expression of tag-fused mdg4 ORFs by immunofluorescent staining. Gag-Flag (green), Pol-StrepII (green), and Env-Myc (green) co-stained with TDP-43-V5 (cyan), H2B-mCherry (red), and DAPI (blue). Scale bar=5 µm. **d**, Images showing effects of mdg4 ORFs expression on the TDP-43 aggregation, co-stained with TDP-43-V5 (green), H2B-mCherry (red), and DAPI (blue). Scale bar=5 µm. **e**, Quantification of cell percentage with cytosolic TDP-43-V5 aggregates after each mdg4 ORF expression. (H2B-mCherry 48.59±2.38%, n=37; Gag-Flag 40.01±3.91%, n=25; Pol-StrepII 49.25±3.15%, n=20 and Env-Myc 47.72±2.83%, n=20). Data are shown as mean ± SEM and a one-way ANOVA with multiple comparisons was performed. ns, no significant difference. **f**, Illustration of the characterization process to test the interaction between pTDP-43 protein and mdg4 RNA by RIP. **g**, RT-PCR result from f to show the mdg4 RNA existing in the TDP-43 aggregate complex.

**Extended Data Fig. 13 Prediction of UG-repeat regions and the interactions between separate R elements on HERV-K RNA. a**, Schematic of potential TDP-43 binding regions on HERV-K RNA. **b**, Computational prediction of potential R-R interacting regions in HERV-K RNA. **c**, Multiple sequence alignment of LTRs across HERV families. **d**, A phylogenetic tree based on LTR sequences across HERV families.

