## Supplementary figures and images for "Human Endogenous Retrovirus K RNA triggers cytoplasmic TDP-43 aggregation by initiating pathological nuclear condensates"

### Supplemental Figure 1

Extended Data Figure 1

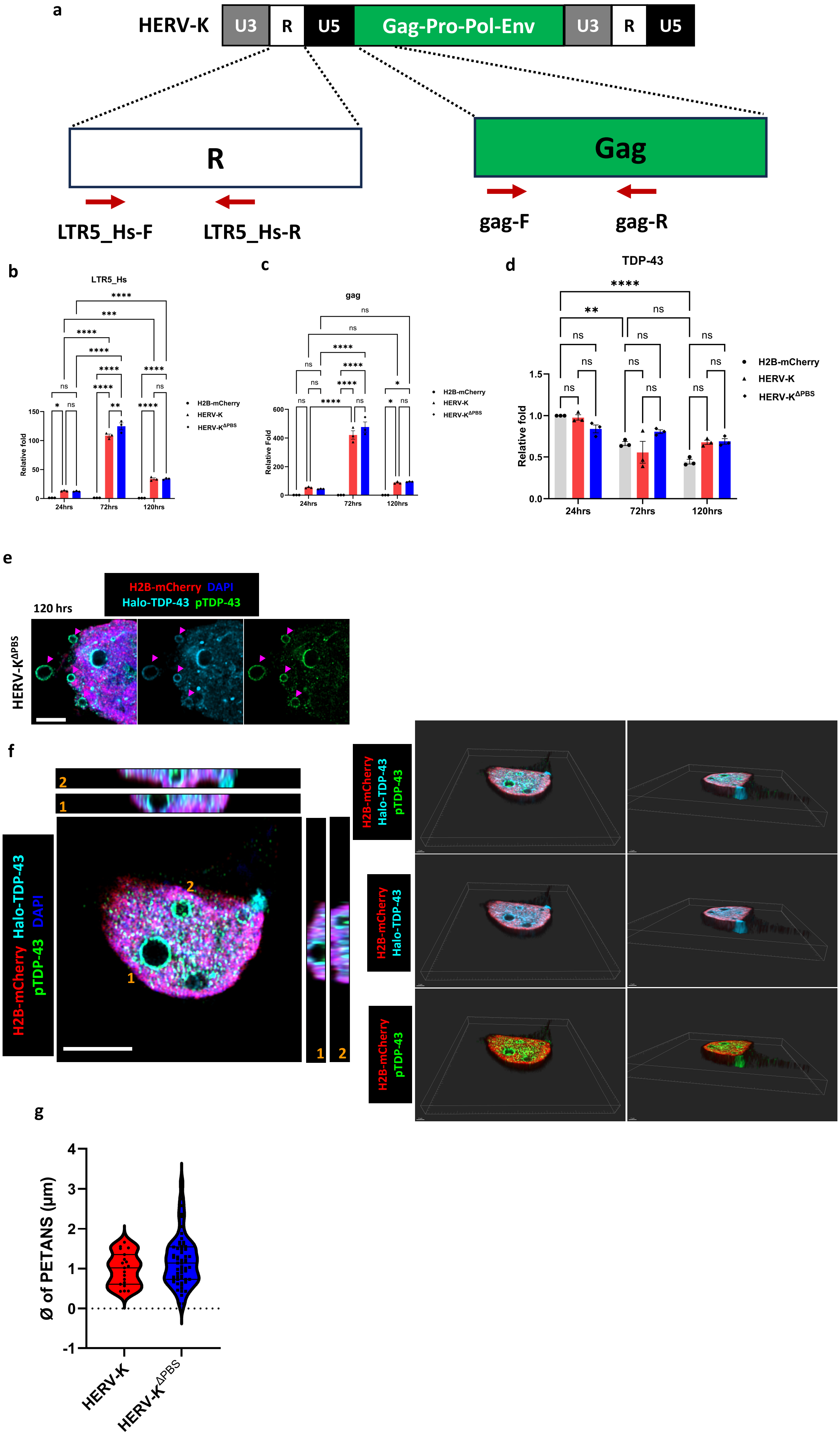

### Supplemental Figure 2

Extended Data Figure 2

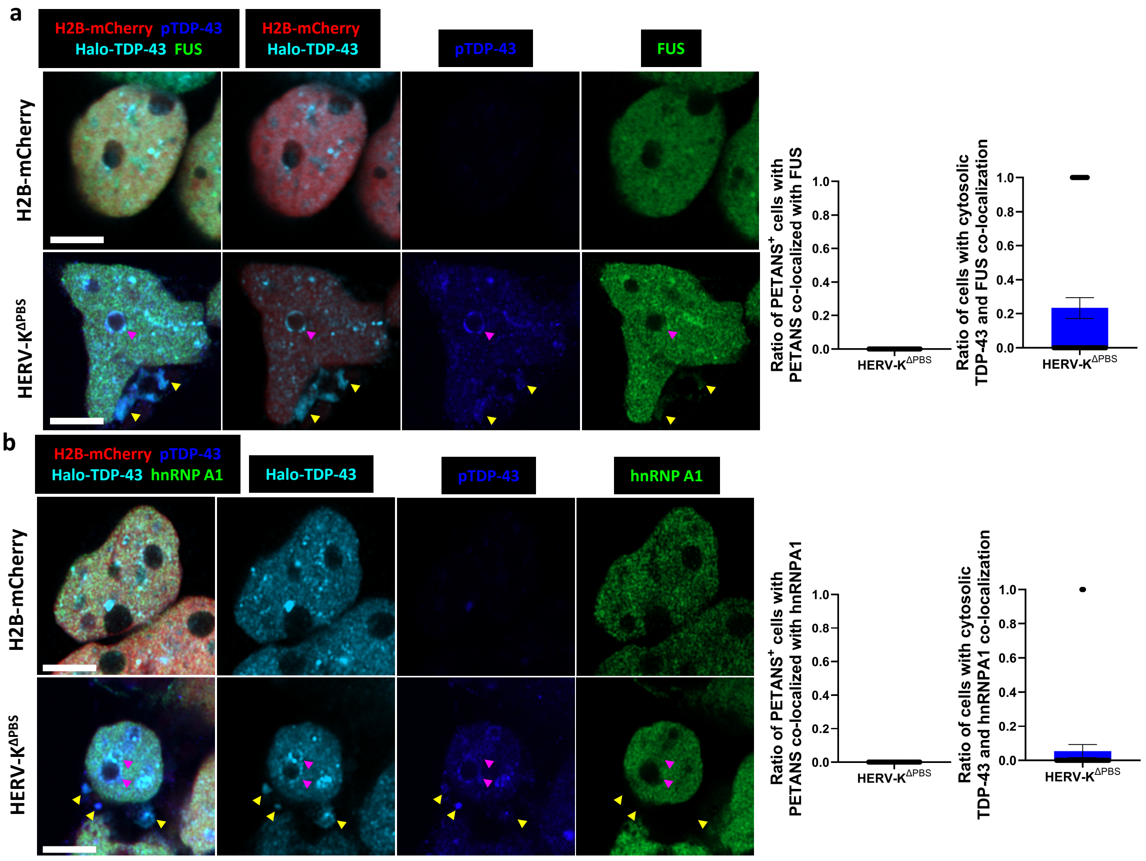

### Supplemental Figure 3

Extended Data Figure 3

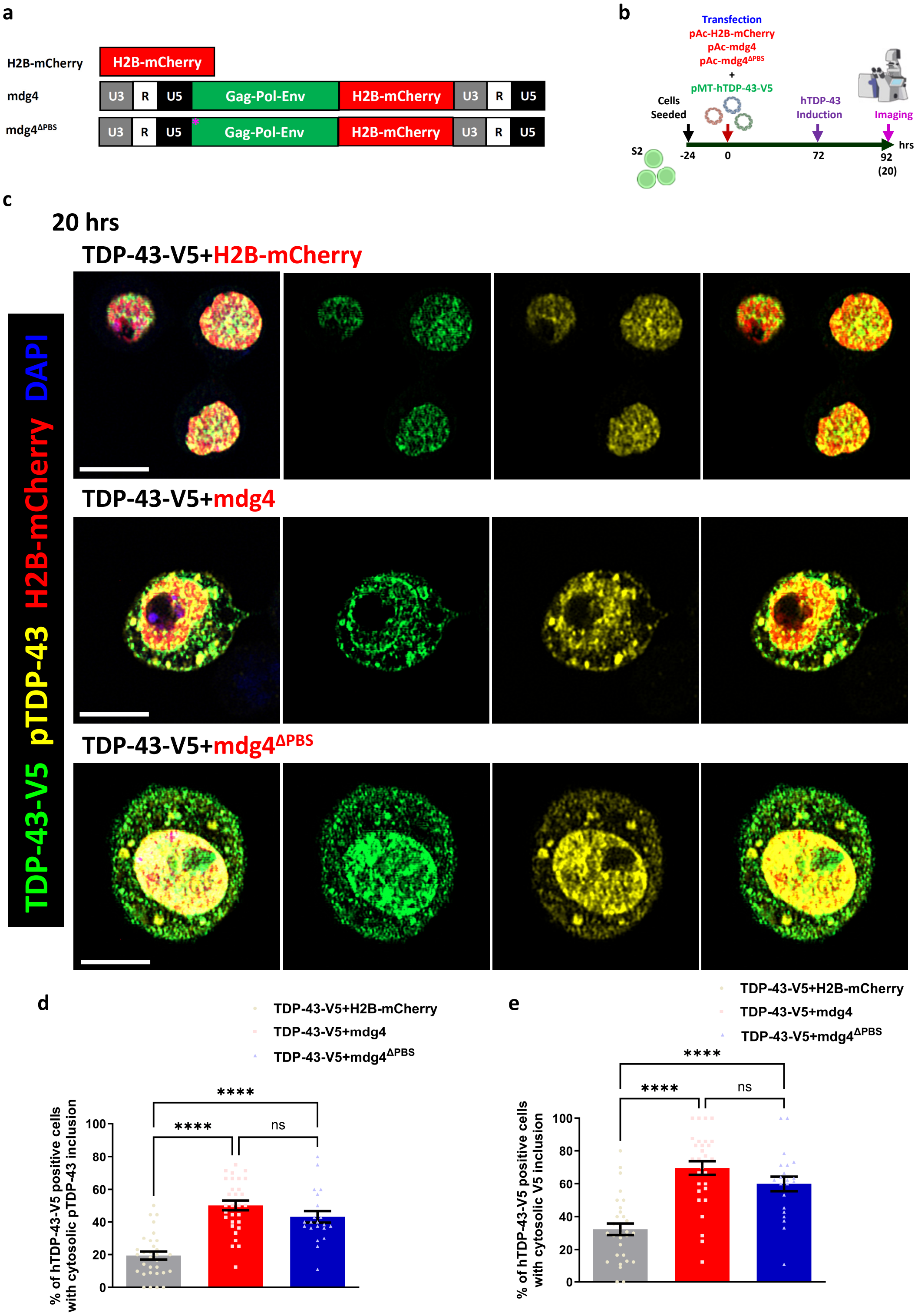

### Supplemental Figure 4

**Extended Data Figure 4**

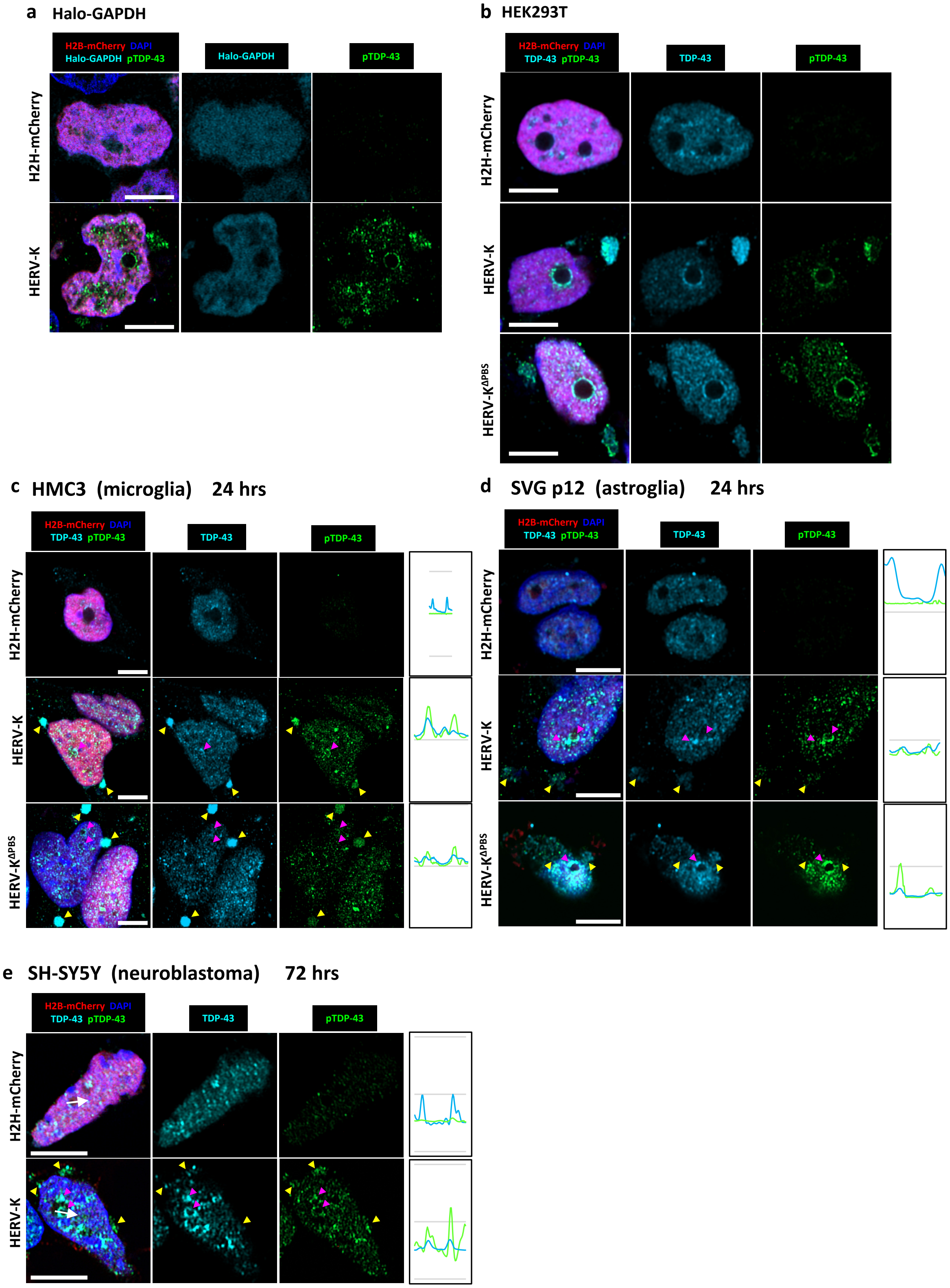

### Supplemental Figure 5

Extended Data Figure 5

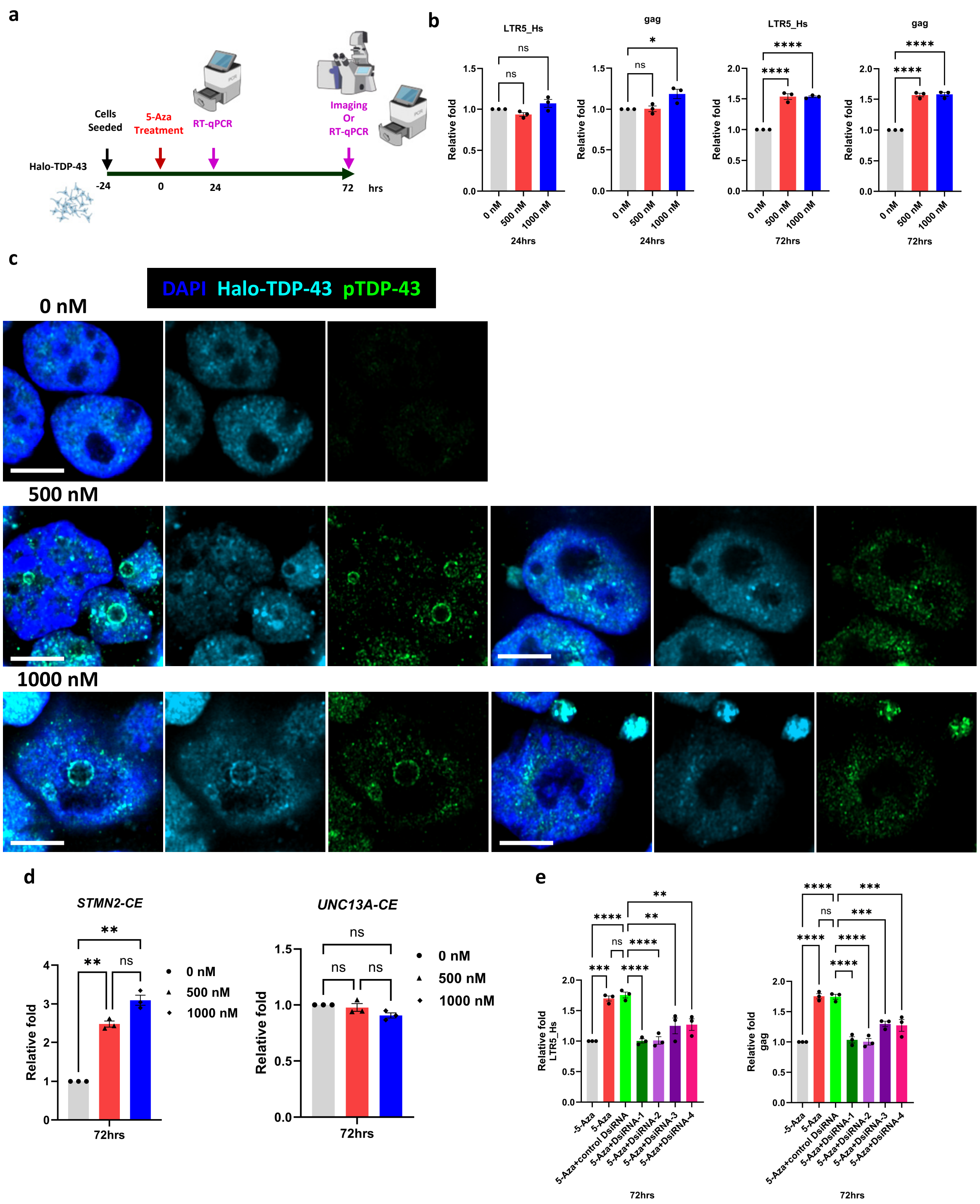

### Supplemental Figure 6

#### Extended Data Figure 6

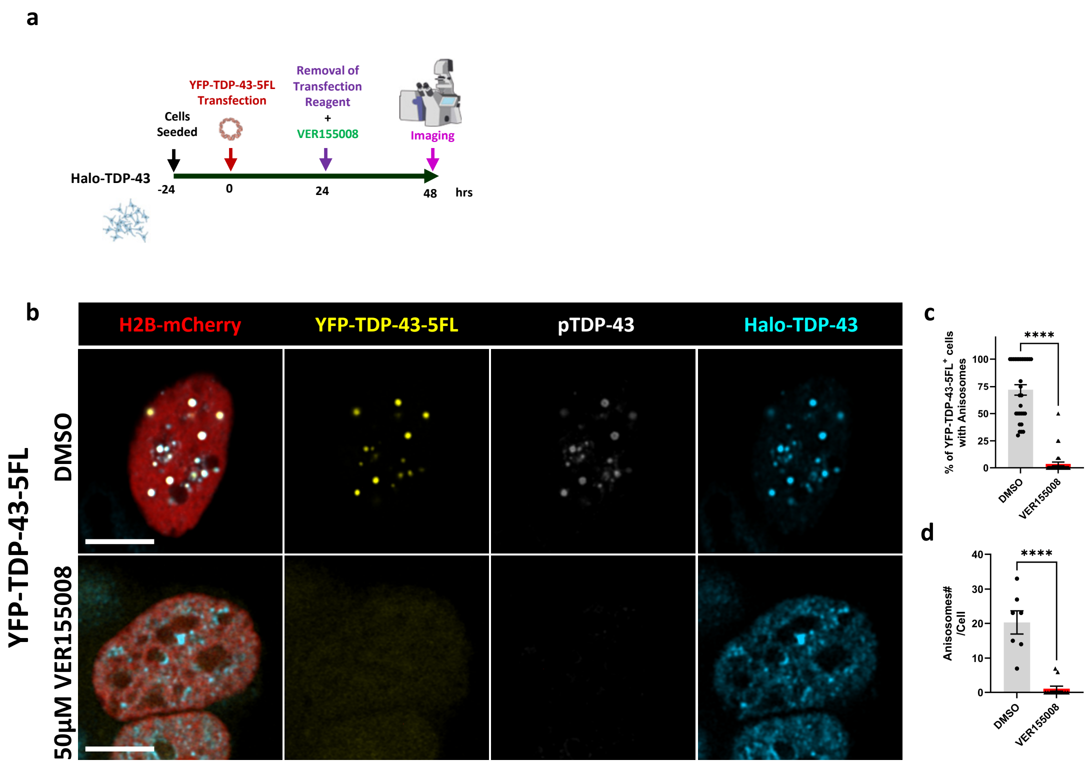

### Supplemental Figure 7

Extended Data Figure 7

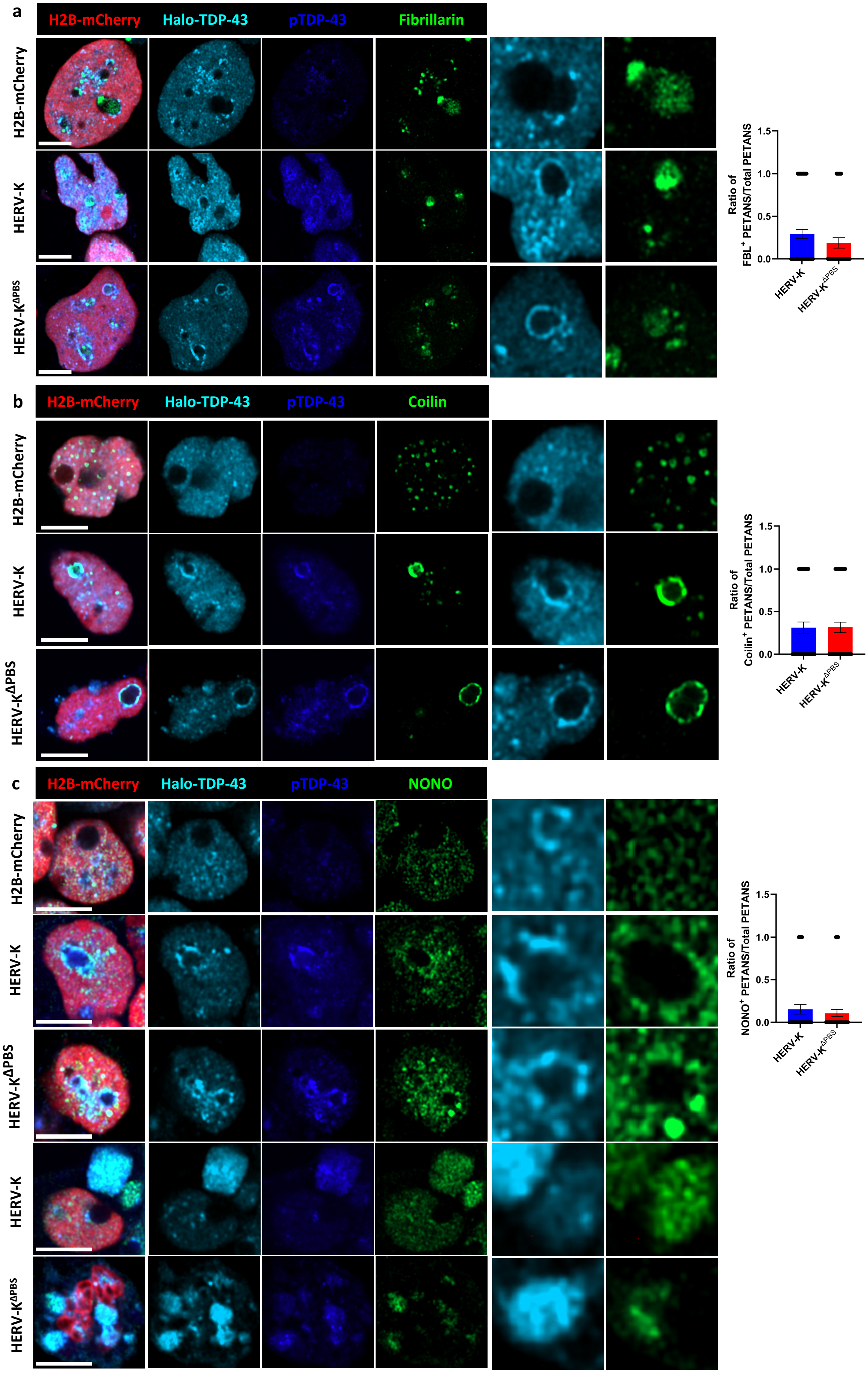

### Supplemental Figure 8

**Extended Data Figure 8**

a

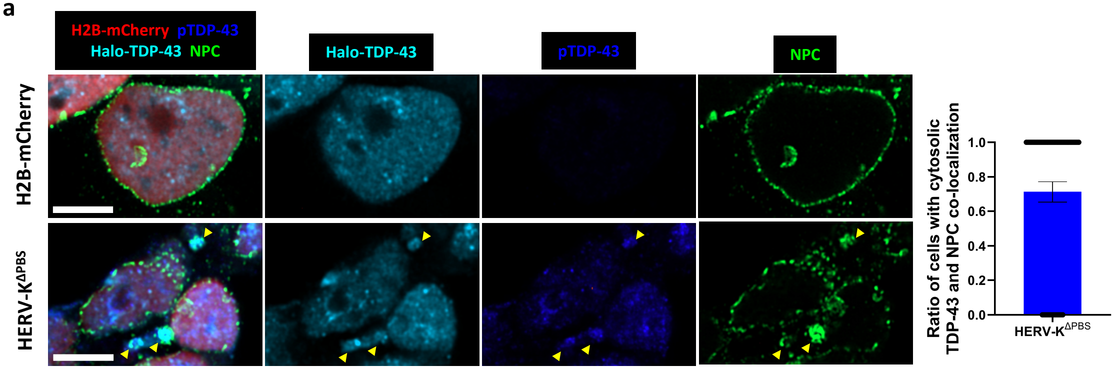


b

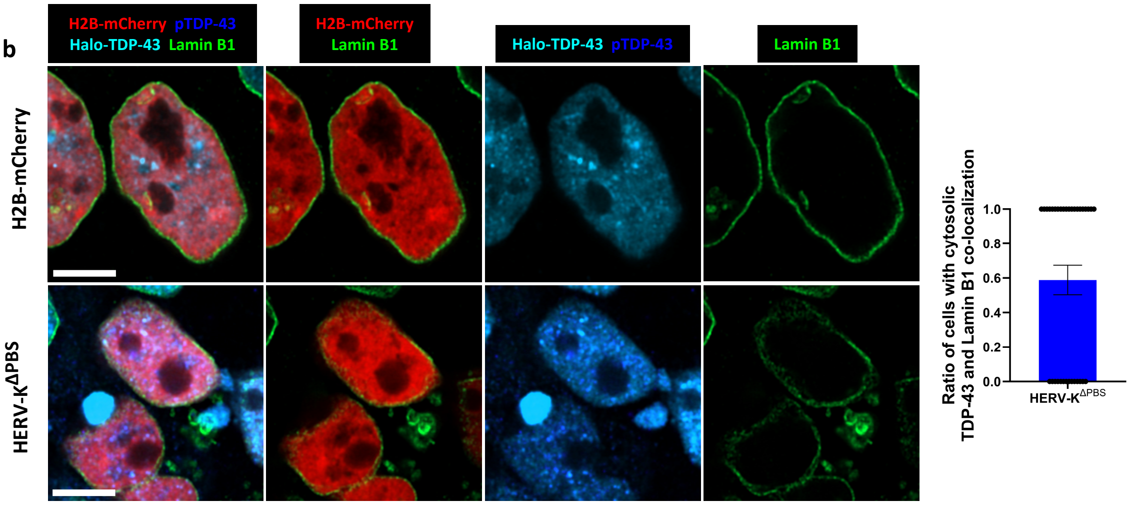

### Supplemental Figure 9

Extended Data Figure 9

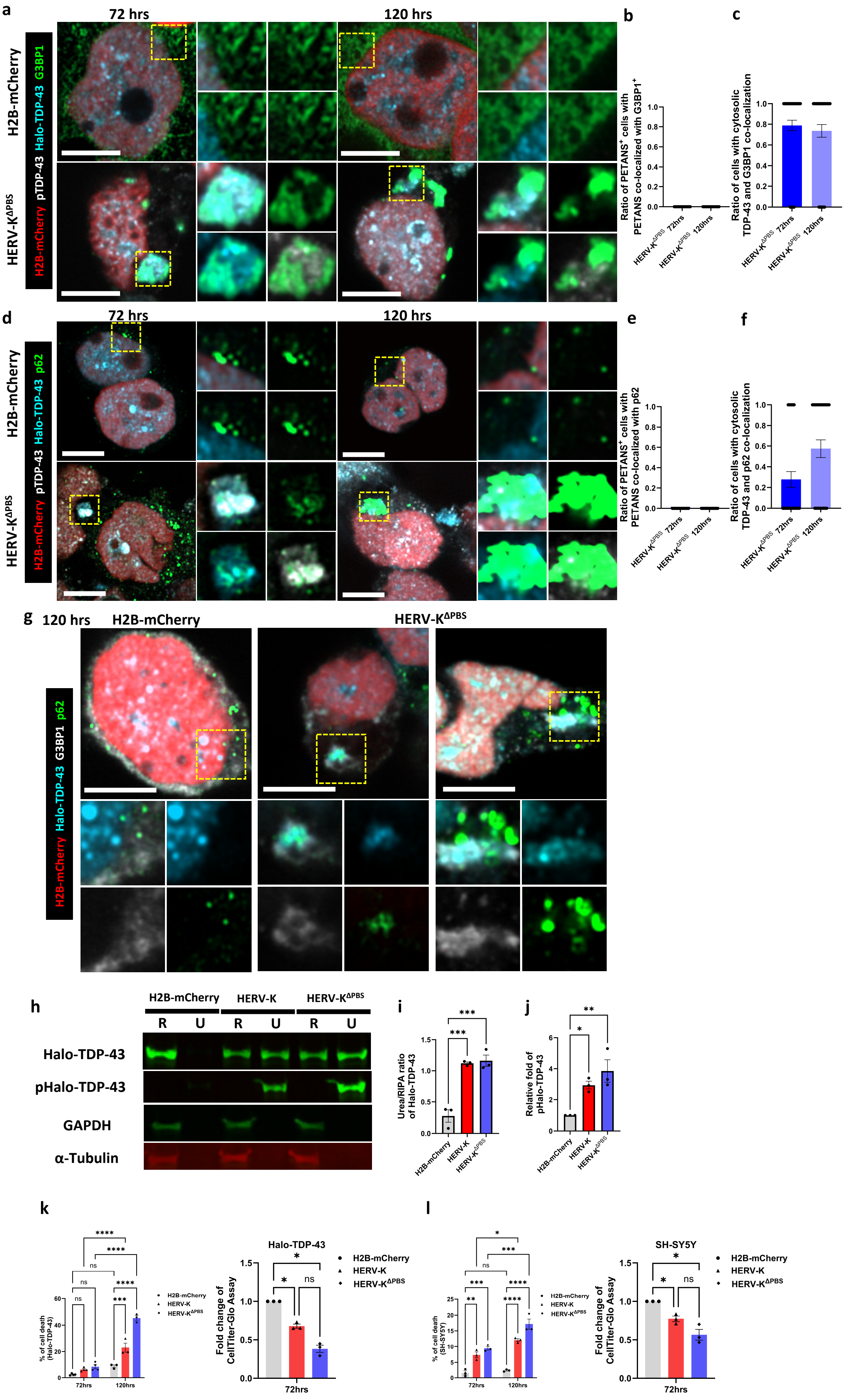

### Supplemental Figure 10

Extended Data Figure 10

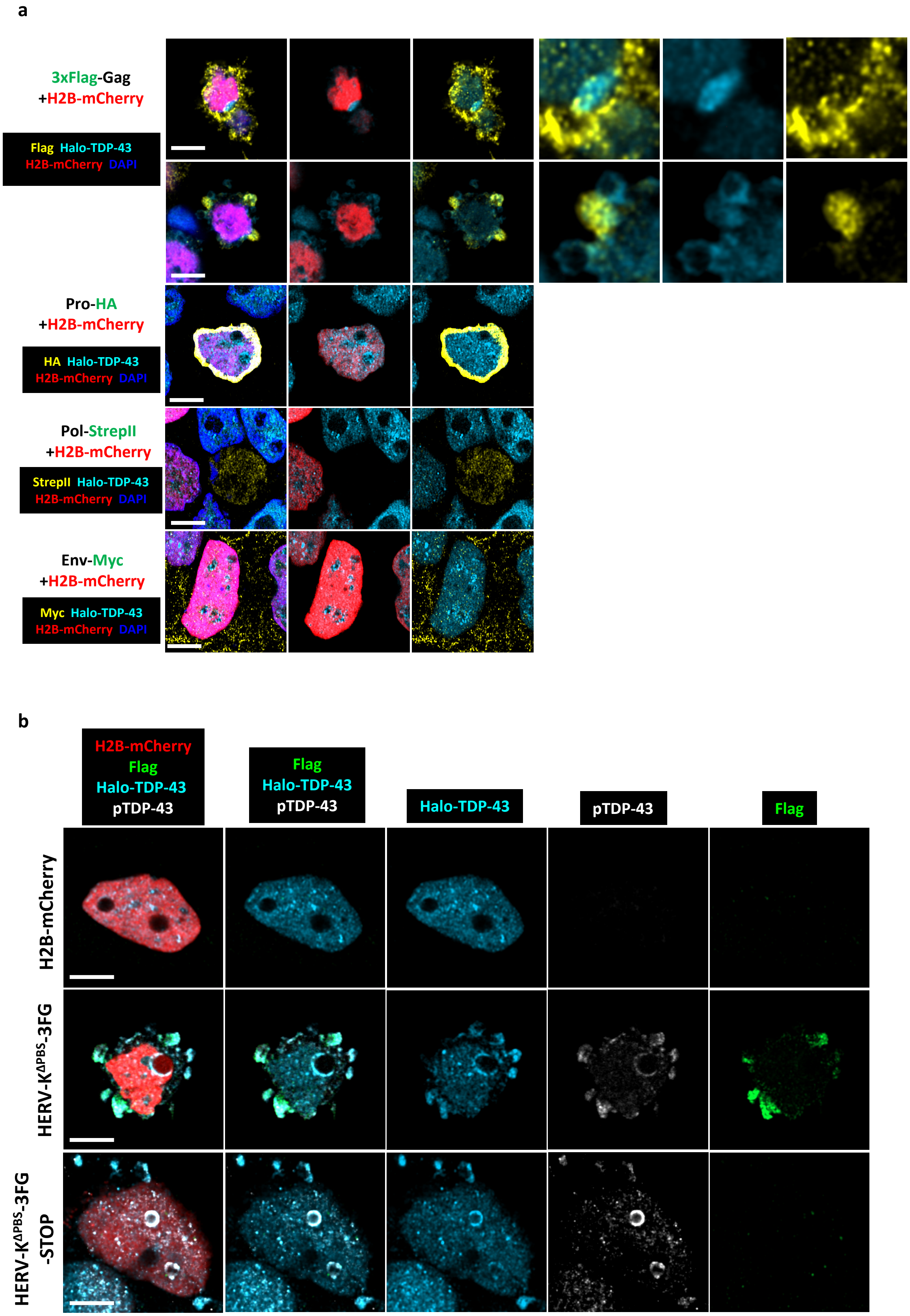

### Supplemental Figure 11

Extended Data Figure 11

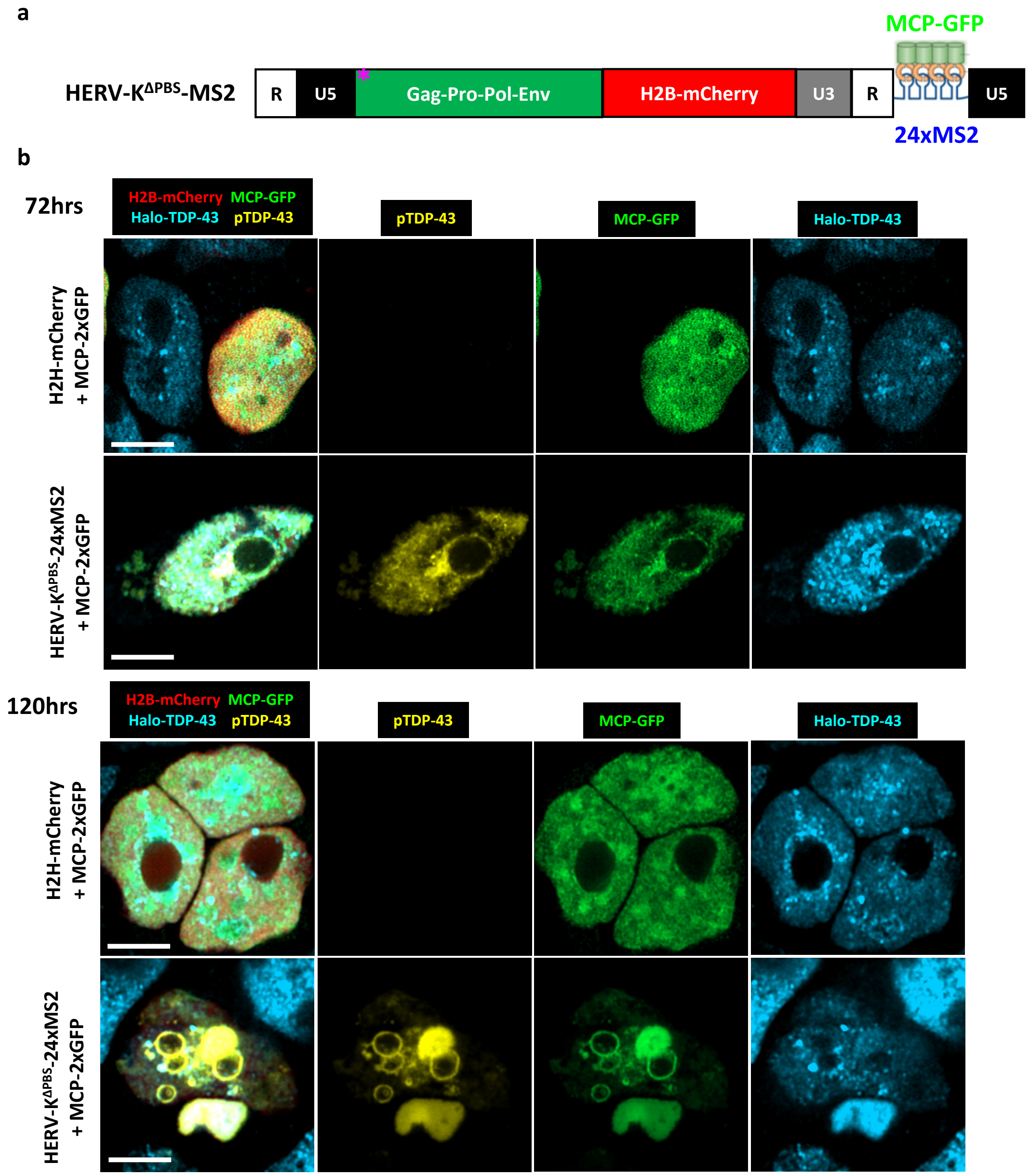

### Supplemental Figure 12

Extended Data Figure 12

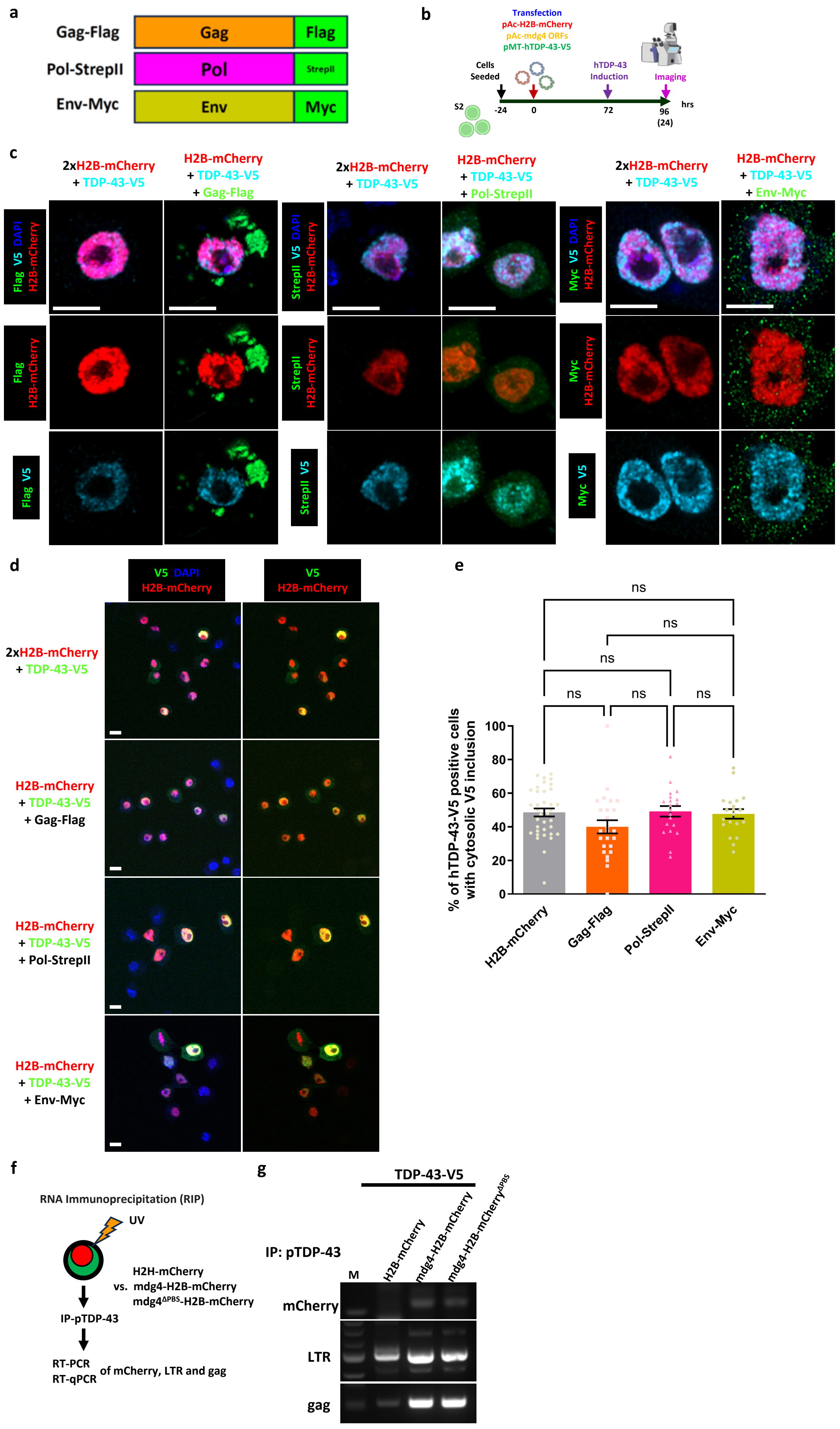
