## Supplemental Figure 13 for "Human Endogenous Retrovirus K RNA triggers cytoplasmic TDP-43 aggregation by initiating pathological nuclear condensates"

**Extended Data Figure 13**

**a**

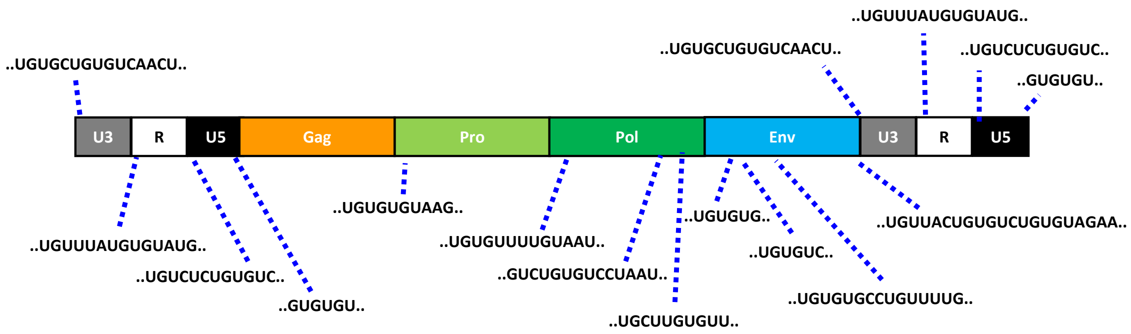


**b**

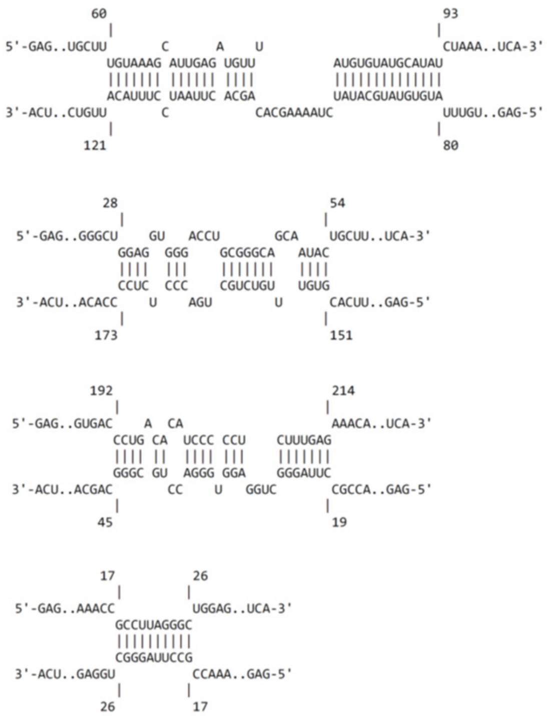


**c**

```
HERV-P(LTR9)        -ttctc--------ctttctcttttcttttgcctattaaac-ctc---------cgctcc 554
HERV-W(LTR17)       -actgg--------gttccacggttctcttccgtgacccac-ggcttctaatagagctat 613
HERV-K(LTR11;HML-8) -aacga-------agcaagaggtctccggaggaagagatga-at-gaat-tagcctatta 582
HuERS-P2(LTR1)      gtccttgcgacaaggcagagggtctaactgagctggttaacacaagccgcctatagatgg 578
HERV-H(LTR7)        ggccgg--------------tccttgccttaagtgatgaca-ttaccttgtaagagtcct 198
HERV-K(LTR3;HML-6)  -cccca-------cgtgcagtgttccttgtggttatctata-ggaatatctggaggctgt 198
HERV-K(LTR13;HML-4) aggagtaccacagtcttcgatctttctgataagtgcataga-agaaacgctgacgtttgc 713
HERV-K(LTR5;HML-2)  ttgtaaagc----attgagatgtttatgtgtatgcatatc-taaaagcacagcatttaat 674
```

**d**

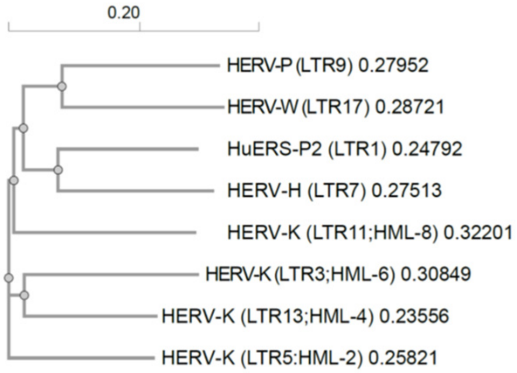
